# Impact of photobleaching of fluorescent proteins on FRET measurements under two-photon excitation

**DOI:** 10.1101/2024.04.24.590958

**Authors:** Dhruba P. Adhikari, Michael R. Stoneman, Valerica Raicu

**Affiliations:** Department of Physics, University of Wisconsin-Milwaukee, Milwaukee, WI 53211, USA

## Abstract

Förster resonance energy transfer (FRET) is a widely used technique for nanoscale molecular distance measurements, which makes FRET ideal for studying protein interactions and quaternary structure of protein complexes. In this work, we were interested in how photobleaching of donor and acceptor molecules affects the FRET results under various excitation conditions. We conducted a systematic study, under two-photon excitation, of the effects of the excitation power and the choice of excitation wavelengths upon the measured FRET efficiencies of multiplex protein constructs, consisting of one donor and either one or two acceptors, using both the kinetic theory of FRET and numerical simulations under given excitation conditions. We found that under low excitation power and properly chosen excitation wavelengths the relationship between the FRET efficiency of a trimeric construct ADA agrees within 2% with the FRET efficiency computed (via the kinetic theory of FRET in the absence of photobleaching) from two dimeric constructs ADN and NDA. By contrast, at higher excitation powers the FRET efficiencies changed significantly, due to the photobleaching of both the donor (through direct excitation) and the acceptor (mostly through FRET-induced excitation). Based on these results and numerical simulations using a simple but powerful algorithm, we also provide guidelines for choosing appropriate experimental conditions for reliable FRET measurements in complexes of associating molecules of biological interest.

## 1. INTRODUCTION

Förster resonance energy transfer (FRET) is a physical phenomenon that involves the non-radiative transfer of energy between an excited fluorescent molecule, called a donor (D), and a neighboring unexcited fluorescent molecule called an acceptor (A) via dipole-dipole interaction [1-3]. FRET has been used to probe the proximity of individual molecules within protein complexes [4, 5], track complex formations [6], monitor dynamic protein interactions [7, 8], and detect conformational changes of macromolecules [9-12] in living organisms. More recently, a FRET technique, called FRET spectrometry, has been developed, which can determine the relative distances between protomers within oligomeric complexes (i.e., quaternary structure) in living cells [13-18].

In order to determine the quaternary structure of biological macromolecules using FRET, it is crucial to understand how the mechanism of FRET works when more than one acceptor are close to an excited donor, as can occur for macromolecules larger than a dimer. Fortunately, the kinetic theory of FRET [19] provides a theoretical basis for predicting the FRET occurring in multimeric complexes. According to the theory, the FRET efficiency of an oligomer containing multiple acceptors can be predicted from the FRET efficiency occurring between each individual donor-acceptor pathway present in the complex. The theory was tested using simulated data [20] and found to accurately predict the FRET occurring in oligomeric complexes in the particular situation where proximity FRET [20] is minimal. The first experimental attempt to test the kinetic theory of FRET [21] found an ∼15% discrepancy between the measured FRET efficiency of a multimeric complex consisting of a single donor and three acceptors vs. the FRET efficiency predicted by the kinetic theory. A separate investigation of the same constructs found a substantially smaller discrepancy (4%) but still concluded the discrepancy was statistically significant [22].

A recently published theoretical analysis of the FRET equations identified several assumptions traditionally used to compute FRET efficiency from experimental data that could lead to significant systematic errors if not carefully considered within the context of the experimental protocols used [23]. Improperly applied assumptions could explain some of the discrepancies between measured and theoretically predicted FRET values, which were observed in experiments testing the kinetic theory of FRET. For instance, in FRET measurements performed using continuous wave (cw) excitation, the equations for determining the FRET efficiency depend on the ratio between the probability of donors to be in the ground state in the presence and absence of FRET; these probabilities have typically been set equal to one in traditional FRET applications, even though when using cw excitation, the value is significantly different than one.

The error caused by the dependence on the integrated probability can be avoided by using pulsed lasers, as it is done in systems relying on two-photon excitation. For two photon excitation of, any discrepancies between measured and theoretical FRET efficiency values are due to a general inability to assess the contributions of photophysical effects, such as photobleaching [24, 25] and photoconversion [25-27] of donors and acceptors, to the measured FRET efficiencies for chosen laser excitation powers.

In addition, since most fluorescent proteins present short Stokes shifts the excitation wavelengths available to excite the donors quite often also lead to direct excitation of the acceptor, which results in an erroneous FRET efficiency readout. Contributions of direct excitation of the acceptor to the measured FRET efficiency can be corrected for by scanning the sample of interest at two [14, 23], or sometimes three [28] different excitation wavelengths instead of one. This in turn may result in additional, caused by the additional excitation wavelengths, which are typically not taken into account but could also affect the FRET efficiency calculation as these effects inevitably change the proportion of donors and acceptors actively involved in FRET.

To avoid significant errors in FRET spectrometry studies and thus accurately determine the interprotomeric distances within oligomers, it is critically important to map out the contributions of photophysical effects to experimentally determined FRET efficiencies under a range of experimental conditions. In this work, we performed two-photon (2P) micro-spectroscopy measurements on living cells expressing FRET constructs of the type used in previous studies [22, 29], consisting of single donor molecule linked to either its C- or N-terminus with one acceptor to form dimers, or linked to both termini with acceptors and therefore form FRET trimers. The kinetic theory of FRET was used to determine the range of excitation powers and combination of wavelengths for which correct relationship exists between the FRET efficiency of dimers and those of trimers. In addition, for comparatively higher excitation powers, we used a simple but effective numerical method to estimate the degree to which photobleaching of donors and acceptors was responsible for the observed discrepancies between the two sets of FRET efficiencies. We quantified photobleaching effects and the extent of their contributions to FRET measurements, as well as identified the general conditions necessary for performing accurate FRET spectrometry measurements using fusion fluorescent proteins [14-17].

## 2. MATERIALS AND METHODS

### 2.1 Fluorescent protein constructs

The plasmids for all the protein constructs used in this work are gifts from Dr. Stephen Vogel. They were expressed and localized in the cytoplasm of Chinese hamster ovary (CHO) cells (see section 2.2). The fluorescent protein fusion constructs [29] used to study FRET were made from combinations of three domains (see Fig. 1): Cerulean [30], as a donor (D) of energy, Venus [31], as an acceptor (A) of energy, and Amber, a non-fluorescent (N) structural place holder containing a point mutation preventing chromophore formation. Two of the constructs (denoted as NDA and ADN hereafter) are dimeric from the FRET standpoint, as they consisted of a single D and a single A involved in energy transfer. The third construct (denoted as ADA) is referred to below as a trimer, since it contains three fluorescent domains, consisting of two As and one D. Cells expressing a construct consisting of Amber fused to Cerulean [29] (i.e., Amber-5-Cerulean, A5C) were also imaged to obtain the emission spectrum for Cerulean. Finally, the emission spectrum of the acceptor was acquired by imaging CHO cells expressing the Venus protein [31], which is also localized in the cytoplasm.

**Figure 1.**
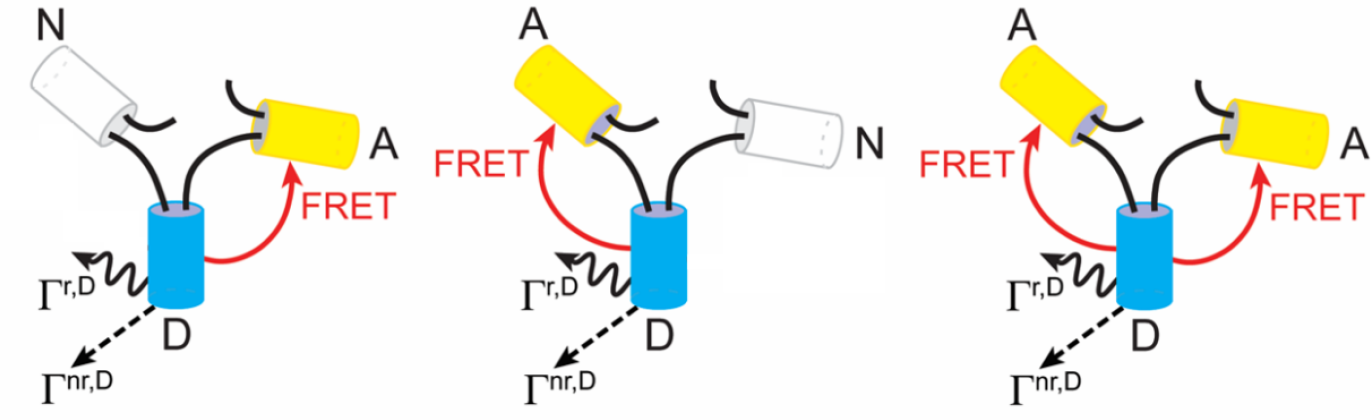
Schematic representation of cytoplasmic FRET constructs and the various pathways for energy loss from the excited donor. White cylinders represent amber (a non-fluorescent structural placeholder, N), cyan cylinders represent the fluorescent molecule Cerulean (a donor of energy, D), and yellow cylinders represent Venus (an acceptor of energy, A). The various arrows indicate possible pathways through which the excited donor loses energy. Wavy arrows indicate radiative loss (Γ^r,D^), dashed arrows indicate non-radiative loss (Γ^nr,D^), and solid arrows pointing toward A from D depict energy transfer through FRET (Γ^FRET^).

### 2.2 Cell culture and protein expression through transient transfection

CHO cells were grown in Dulbecco’s modified Eagle’s medium (DMEM) supplemented with 10% fetal bovine serum, 1% L-glutamine, 1% penicillin/ streptomycin, and 1% non-essential amino acids. All reagents were purchased from Fisher Scientific unless noted otherwise. Cells were seeded onto 35-mm No. 1.5 coverglass culture dishes (Part No. NC079415, Cellvis, CA), at a density of 7 × 10^4^ cells per dish; the dishes incubated for 48 h at 37 °C in a humidified environment with 5% CO_2_. One such dish was prepared for each of the samples mentioned in section 2.1, for a total of 5 dishes. After incubation, each dish was transfected with a single plasmid encoding for one of the five constructs described above. A total of 2 μg plasmid DNA was used for the transfection procedure, along with 8 μl lipofectamine 2000 (Invitrogen, Carlsbad, CA) and diluted in 250 μl OptiMEM (Invitrogen), according to a previously described procedure [32]. After transfection, the dishes were returned to the incubator. After 24 h, the dishes were removed from the incubator and taken for imaging. For imaging, the cell growth medium was removed, and the cells were first rinsed with Dulbecco’s phosphate-buffered saline (DPBS; REF:14190-144, Life Technologies, NY) then resuspended in DPBS.

### 2.3 Spectrally resolved imaging

Spectrally resolved fluorescence images of CHO cells expressing the cytoplasmic fluorescent constructs were acquired using a previously described two-photon (2P) optical micro-spectroscope [33]. This imaging system consisted of a mode-locked Ti-Sapphire laser (Mai Tai™, Spectra Physics, Santa Clara, CA), which generated 100 fs pulses with tunable central wavelengths between 690 nm and 1040 nm and was used as the excitation source, an OptiMiS™ scanning/detection head (Aurora Spectral Technologies, Grafton, WI), and an inverted Nikon Eclipse Ti™ microscope (Nikon Instruments, Inc. Melville, NY) equipped with an infinity-corrected oil-immersion objective (NA=1.45, 100×). The OptiMiS detection head used a non-de-scanned acquisition scheme in which emitted fluorescence was spectrally resolved by passing it through a transmission grating and projecting it onto a cooled electron-multiplying charge-coupled device (EMCCD) camera (iXon Ultra 897; Andor Technologies, South Windsor, CT), similar to previously described systems [18, 34]. The excitation beam was focused to a diffraction-limited spot which was scanned across the sample along an entire line consisting of 440 pixels, with a pixel dwell time of 35 milliseconds (i.e., 80 μs pixel dwell time) while the camera integrated the fluorescence signal during the course of the entire sweep. Three hundred such lines were scanned, which each line providing an entire emission spectrum for each pixel. By raster scanning the beam across the sample in this manner, we acquired 3D fluorescence micro-spectroscopic image stacks, with two dimensions representing spatial distribution of the fluorophores and the third dimension corresponding to the emission spectrum. The spectral resolution was set at 1.1 nm.

Three different pairs of excitation wavelengths (800 nm/880 nm, 800 nm/960 nm, and 880 nm/960 nm, maintaining their order) and various excitation powers (15 mW/point, 42 mW/point, 52 mW/point, and 62 mW/point) were used to explore the impact of excitation wavelength and power on the determination of FRET efficiency. Each field of view was scanned using a single excitation power and a single pair of excitation wavelengths. For each excitation power, four to five experiments were conducted, each of which generated a complete set of measurements of all FRET constructs, acquired using all three pairs of excitation wavelengths (∼10 fields of view, or about 30 cells, per excitation power per wavelength pair). All measurements were taken by focusing the laser on the cross-sections of CHO cells, particularly the parts corresponding to the cytoplasm, which were cultured on the coverslip surface as described in section 2.2.

### 2.4 Image analysis and calculation of FRET efficiency

#### 2.4.1. Spectral unmixing and segmentation of intensity map

The composite emission spectrum from each pixel in the micro-spectroscopic images of cells expressing cytoplasmic trimeric FRET constructs was deconvolved (unmixed) into donor (D) and acceptor (A) components using linear regression, as described elsewhere [14, 33, 35]. The elementary spectra for Cerulean [30] (donor) and Venus [31] (acceptor) used in the unmixing procedure were obtained by imaging cells expressing A5C only and Venus only, respectively [22].

By applying the unmixing procedure to each pixel of a fluorescence micro-spectrograph, we obtained 2D maps of donor intensity in the presence of acceptor, *k*^*DA*^, and acceptor intensity in the presence of the donor, *k*^*AD*^. The *k*^*DA*^ and *k*^*AD*^ values thus obtained were multiplied by the area underneath the elementary spectra of D and A, respectively, to obtain the total donor fluorescence emission in the presence of the acceptor, *F*^*DA*^, and total acceptor fluorescence emission in the presence of donor, *F*^*AD*^, for each image pixel. Pixel-level apparent FRET efficiency values, *E*_*app*_, were calculated using the corresponding *F*^*DA*^ and *F*^*AD*^ at each pixel (see section 2.4.2). *F*^*DA*^ map were used to manually demarcate regions of interest (ROIs) in the micro-spectroscopic scans, which were then segmented into areas of ∼900 pixels using a moving-square algorithm described in a previous publication [36]. The pixel locations for each segment were saved and paired to the corresponding image source. In the dual-wavelength excitation approach, segment-level FRET efficiency values were calculated using the average *F*^*DA*^ and *F*^*AD*^ obtained for the corresponding segment (see section 2.4.3).

#### 2.4.2. *E*_*app*_ *calculation using single-wavelength excitation*

For excitation wavelengths that do not lead to appreciable direct excitation of the acceptor, the FRET efficiency can be calculated from using the following equation [33]:

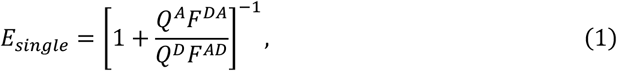

where *Q*^*D*^ and *Q*^*A*^ are the quantum yields of the donor (D) and acceptor (A), respectively. Determining FRET efficiency values using the single-wavelength excitation method is simpler to implement and less prone to calibration errors but it could lead to inaccurate FRET efficiency values when its underlying approximation does not hold.

Following, pixel-level *E*_*single*_ calculation, we split the ROIs in each cell into segments, as described above, and a histogram of *E*_*single*_ values was constructed for each segment using a bin size of 0.005. Then, the dominant peak of each individual segment histogram was extracted and used to create a “meta-histogram” of the dominant peak positions. The bin size used to construct the meta-histogram was chosen to be equal to 0.02, as described previously [15, 17, 18]. Finally, we extracted a measured value of the FRET efficiency, 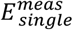, from the meta-histogram for each FRET construct using the dominant peak (mode) position of the meta-histogram.

#### 2.4.3. E_app_ calculation using dual wavelength excitation

Scanning the sample at two excitation wavelengths allowed us to obtain the concentrations of both Ds and As, and to correct the computed FRET efficiency from inadvertent contributions from direct excitation of the acceptors. Spectral unmixing of such images resulted in four separate 2D maps: the fluorescence signal of the donor obtained from the first 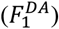 and second 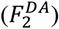 excitation wavelengths and the acceptor emission at the first 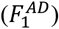 and second 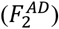 excitation wavelengths. In this case, the FRET efficiency was computed via the following equation [23]:

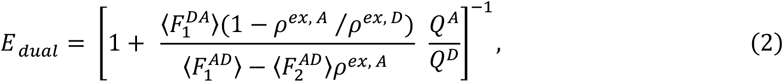

where *Q*^*D*^ and *Q*^*A*^ are the quantum yields of the donor and acceptor, respectively, the subscript in 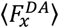 or 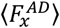 is *x* = 1 for the first excitation wavelength, and *x =* 2 for the second; while the ⟨ ⟩ symbol represents the average over all pixels in an ROI segment. The quantities *ρ*^*ex D*^ and *ρ*^*ex A*^ are the intensity ratio of the donors and acceptors, respectively, between the two excitation wavelengths, i.e., 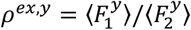 where *y=D or A;* these ratios were obtained by scanning samples expressing only the donor or only the acceptor using the same pair of excitation wavelengths. The dual-wavelength excitation scheme compensates for any direct excitation of the acceptor and also provides the donor and acceptor concentration – which enables the straightforward calculation of the acceptor molar fraction, as described in detail in previous publication [18] and summarized in the Supplementary Method section SM1.

In this study, we used three different excitation wavelength pairs for the dual-wavelength excitation protocol: 800 nm/880 nm, 800 nm/960 nm, and 880 nm/960 nm. The values of *ρ*^*ex A*^ and *ρ*^*ex D*^ for each pair of excitation wavelengths were determined separately by scanning the cells expressing only Venus and only A5C between two excitation wavelengths for each excitation power. The values of *ρ*^*ex A*^ and *ρ*^*ex D*^ for all the excitation wavelength pairs and powers are given in Supplementary Table 1. Using these excitation ratios, a single value of *E* _*dual*_ was calculated per ROI segment (as opposed to per pixel as described in the previous section) using the average values of 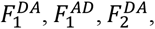, and 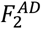 from the corresponding segment, because molecules can diffuse into and out of a pixel within the time it takes to change the excitation wavelength of the laser and scan the sample again. Subsequently, we determined the measured FRET efficiency for the dual-wavelength excitation method 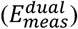 by averaging *E*_*dual*_ values across all segments from all measurements for each specific construct.

**Table 1.** Apparent FRET efficiencies (weighted averages ± weighted standard deviation, SD_w_, or propagated error, δ) for cytoplasmic FRET constructs measured using single-wavelength excitation at 800 nm and with an average laser power of 15 mW/pixel. The final error computed by combining the “measured” and “predicted” errors in quadrature is shown in parenthesis.

| Constructs | Apparent FRET efficiencies |  |  |
| --- | --- | --- | --- |
| | $E_{single}^{meas} \pm SD_w$ | $E_{ADA}^{pred} \pm \delta E_{ADA}^{pred}$ | $\Delta E_{ADA} \pm \delta \Delta E_{ADA}$ |
| NDA | $0.581 \pm 0.023$ | — | — |
| ADN | $0.628 \pm 0.022$ | — | — |
| ADA | $0.781 \pm 0.040$ | $0.754 \pm 0.017$ | $0.027 \pm 0.057$ (0.043) |

#### 2.4.4. Cross-checking the results using the kinetic theory of FRET

The FRET efficiency for the trimeric FRET construct was determined both from direct measurements of said construct and computed from the measurements of the dimeric FRET efficiencies via a formula derived previously (see Supplementary Methods section SM2) from the kinetic theory of FRET [19, 23].

### 2.5 Numerical simulations for estimating the degree of D and A photobleaching

To estimate the effect of D and A photobleaching on the FRET efficiency we used a numerical simulations process (see Supplementary Methods Section SM4) that started with dispersing the trimeric FRET constructs (ADA) randomly onto an array of 1,200 pixels, with 20 such constructs assigned to each pixel. Then, the construct-level fluorescence signals of donor and acceptor were computed for all possible forms of the construct (see the table at the top of Supplementary Fig. SM1). These construct-level fluorescence signals were determined from known absorption coefficients, quantum yields, and the measured FRET values of the constructs (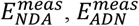, and 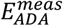) obtained at the lowest excitation power of 15 mW (i.e., the power at which photobleaching was minimal).

Next, individual donors and acceptors in the pixel array were randomly “turned off” (i.e., photobleached), so that they do not participate in FRET. The decision process for photobleaching relied on one of the inputs photobleaching parameters. For the first step, the parameters needed were the probabilities that a donor or acceptor undergoes photobleaching after the excitation at *λ*1 was completed, i.e., 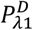 and 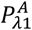, respectively. If the fluorescent molecule was assumed as photobleached, it was then converted into a non-fluorescent molecule, N, and the original ADA construct was reclassified as a different species. If, for example the random decision process resulted in the first A in an ADA construct being photobleached while the other two molecules remained unbleached, the resulting construct after this step became NDA. If the donor was assumed as photobleached the same construct became ANA, which is not FRET productive.

After repeating the process of assigning photobleached (i.e., N) status to a fraction of the molecules in each simulated pixel, we computed the total donor fluorescence signal 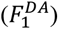 and acceptor fluorescence signal 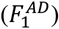 by summing up the emitted signal from each construct in said pixel. This is akin to collecting the fluorescence intensity values resulting from the 1^st^ excitation wavelength in the measured data. The values of 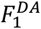 and 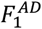 were calculated according to the following formula:

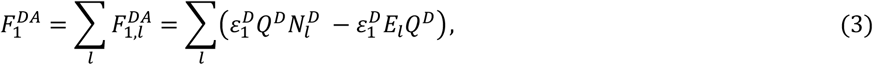

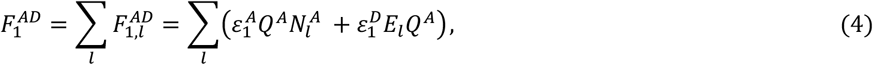

where the summation extends over all the constructs, *l*, in that particular pixel, and 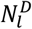 and 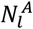 are the number of D and A in a given construct arrangement, *l*, and *E*_*l*_ the FRET efficiency of said construct arrangement. Due to the possibility of undergoing photobleaching, there are *l* = 8 potential arrangements of the ADA construct which could arise due to photobleaching, i.e., ADA, ADN, NDA, NDN, ANA, ANN, NNA and NNN.

Next, we simulated the effect of random photobleaching during the second excitation step, i.e., at the second excitation wavelength. The process was similar to the first simulation step, except for utilizing a different set of input photobleaching parameters, i.e., 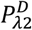 and 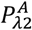. The fluorescence intensities of each pixel were calculated according to the following formula (which is similar to the one used at step 1):

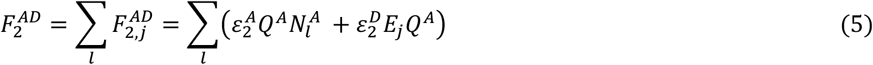

Finally, utilizing the pixel-level fluorescence intensities, we estimated the apparent FRET efficiency for each simulated pixel. We use Eq. (1) to estimate simulated FRET efficiency for the single-wavelength excitation 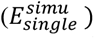 method and Eq. (2) to estimate simulated FRET efficiency for the dual-wavelength excitation 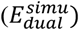 method. After calculating the FRET efficiency in each pixel of the simulated camera array using Eq. (1), we constructed a histogram of 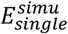 values and extracted the peak of this histogram. We also computed the average 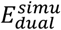 value over the array of pixels using Eq. (2).

As stated above, we employed a 3D array of donor and acceptor photo-bleaching probabilities. Each point in the 3D array contained a unique combination of 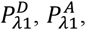, and 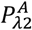. A separate numerical simulation was performed for each point in the 3D array of photo-bleaching probabilities. Furthermore, to reduce potential errors, we repeated the simulation 20 times for each set of photo-bleaching probabilities. For the dual-wavelength method, the value of 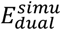 is estimated from the average of the FRET efficiencies obtained from all 20 repetitions. Whereas for single-wavelength method, the dominant peak of histogram from each repetition was extracted and used to create a “meta-histogram” of the dominant peak positions using a bin size of 0.01, similar to what was performed for the measured FRET efficiency values obtained using the single-wavelength method (described in the section 2.4.2). Ultimately, for a single set of photobleaching probabilities, we obtained an average simulated FRET efficiency for both excitation methods, single- and dual-wavelength excitation. This array of simulations was performed for all the FRET constructs, i.e., NDA, ADN, and ADA.

Once the simulated FRET efficiencies were estimated for all three FRET constructs, the predicted FRET efficiency 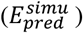 for the trimeric construct was also estimated utilizing simulated FRET efficiencies of dimeric constructs ADN and NDA. Thereafter, the discrepancy 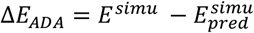 was determined for the trimeric construct, similar to what we have done for the measured data.

After obtaining all the simulated FRET efficiencies for all the constructs for in which the donors and acceptors have been photobleached to various degrees, we systematically compared them to those measured experimentally, by computing fitting residuals using Supplementary Eq. SM16 (see Supplementary Methods section SM4). By identifying the set with the smallest fitting residual relative to the experiment, we de facto identified the degree of photobleaching probability of donors and acceptors at the first scan (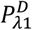 and 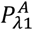, respectively) and photobleaching probability of acceptor at second scan 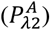.

## 3. RESULTS AND DISCUSSION

### 3.1 Validation of the experimental and analysis protocols under low excitation power

The overall goal of this study is to evaluate the impact of the excitation conditions on the photobleaching of FRET donors and acceptors and how photobleaching affects the accuracy of the results. To do this, we took advantage of important tools available to us, including: (1) Pulsed excitation light (see Materials and Methods) to avoid errors caused by the acceptors and donors being simultaneously in an excited state and therefore unable to be involved in FRET (see Introduction). (2) A trimeric acceptor-donor-acceptor (ADA) construct that presents a high FRET efficiency that is relatively easy to determine with high accuracy, and two additional dimeric constructs, NDA and ADN (where N is a dark place holder) whose FRET efficiencies are used to predict (for comparison to experiment) the FRET efficiency of the ADA construct. (3) An optical micro-spectroscope with high signal-to-noise ratio, which allows us to accurately determine FRET efficiencies at low as well as high excitation powers, i.e., in the presence or absence of photobleaching, respectively.

Two methods have been used in the past to determine the FRET efficiency of oligomeric constructs: (*i*) single- and (*ii*) dual-wavelength excitation schemes. In the first approach, an excitation wavelength is chosen that excites the donor significantly while exciting the acceptor only minimally [23, 33]. This method simplifies the measurement and data analysis process. The dual-excitation method, on the other hand, involves scanning the sample at two different wavelengths. The second excitation scan allows for both the correction of any contribution to FRET by unintended direct excitation of the acceptor at the first excitation wavelength and the estimation of the concentration of donors and acceptors [14, 23]. In this section, we compare the accuracy of these two methods using the lowest excitation power that we could achieve without compromising the signal-to-noise levels.

For the single-wavelength excitation method, we tuned the center wavelength of the laser pulses to 800 nm. This wavelength is known to induce appreciable two-photon excitation of Cerulean but negligible excitation of Venus fluorescent proteins [25]. Typical results for the trimeric construct ADA expressed in the cytoplasm of CHO cells are illustrated in Fig. 2. After spectrally unmixing the composite micro-spectroscopic images of the samples (Fig. 2a and b), the FRET efficiency was computed using Eq. (1) for every pixel to generate two-dimensional FRET efficiency maps (Fig. 2c) on which regions of interest (ROI) were manually drawn and divided into smaller segments (Fig. 2c), as described in Materials and Methods. A histogram was then generated from the *E*_*app*_ values of all pixels within a particular segment (Fig. 2d). Finally, the peak positions of each *E*_*app*_ histogram were collected and used to generate a final distribution of the frequency of occurrence of each peak value, i.e., a meta-histogram of *E*_*app*_ (Fig. 2a). The process was repeated for each FRET construct for all the cell samples investigated. From the peak position of each meta-histogram (Fig. 2e), the weighted average 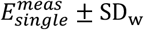 was computed for all the ROIs for each construct using each excitation power.

**Figure 2.**
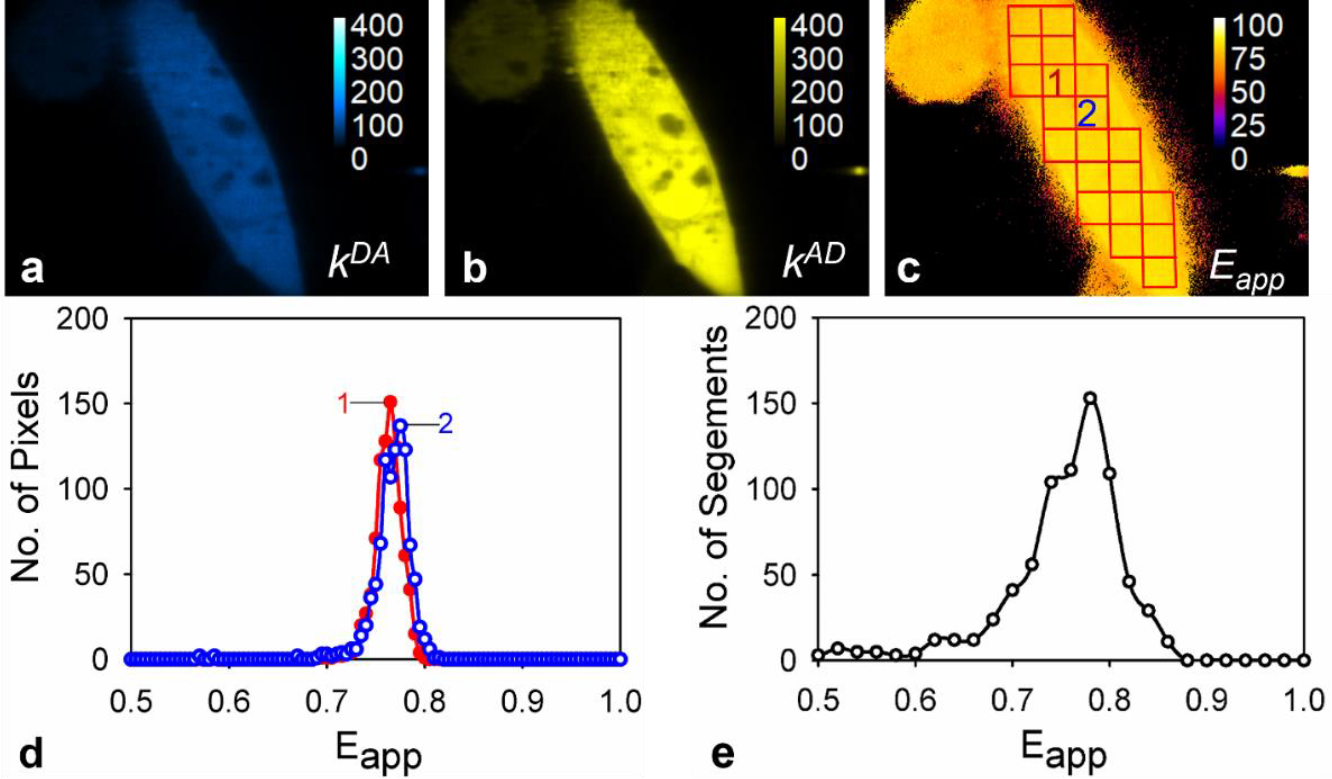
Typical results obtained from imaging CHO cells co-expressing cytoplasmic FRET constructs using two-photon excitation at 800 nm. Unmixing of the original fluorescence micro-spectrographs resulted in the generation of two-dimensional intensity maps for the signal of (a) donors in the presence of acceptors, *k*^*DA*^, and (b) acceptors in the presence of donors, *k*^*AD*^. Apparent FRET efficiency, *E*_*app*_, maps (c) were computed from the pixel-level values of *k*^*AD*^ and *k*^*DA*^. Hand-drawn regions of interest for the *E*_*app*_, maps were divided into smaller segments (red squares) using the *moving-squares* method (see Materials and Methods). Panel (d) shows individual *E*_*app*_ histograms generated (using a bin size of 0.005) for the randomly selected segments labeled with 1 (red line, filled circles) and 2 (blue line, empty circles) in c. A meta-histogram (e) obtained for the FRET construct ADA from a typical experimental dataset (∼ 20 images) was generated by combining the peak positions of individual *E*_*app*_ histograms generated from each ROI segment of all the cells imaged expressing this construct.

Average FRET efficiencies obtained when using single-wavelength excitation for each of the three FRET constructs at the lowest excitation power used in this work (15 mW per pixel) are listed in Table 1 and for all four excitation powers in Supplementary Tables 2-4. As it will become apparent momentarily, the FRET efficiencies obtained for the 15 mW laser power provide good reference points for the subsequent assays, in that they are minimally affected by photobleaching. We note for now that difference between the FRET efficiency value measured for the ADA fusion protein stands in good agreement with the one computed, or “predicted,” from measurements of the NDA and ADN constructs using a relationship obtained from the kinetic theory of FRET (see Ref. [22] and Supplementary Method section SM2). The overall difference is 0.027 or about 3.5%, which is within the standard deviation for these measurements.

**Table 2.** Apparent FRET efficiencies (weighted averages ± weighted standard deviation, SD_w_, or propagated error, δ) for cytoplasmic FRET constructs measured using the two-wavelength excitation protocol for three different pairs of wavelengths average laser powers of 15 mW/pixel for all the wavelengths. The final error computed by combining the “measured” and “predicted” errors in quadrature is shown in parenthesis.

| Constructs | Wavelength pairs (nm/nm) | Apparent FRET efficiencies |  |  |
| --- | --- | --- | --- | --- |
| | | $E_{dual}^{meas} \pm SD_w$ | $E_{ADA}^{pred} \pm \delta E_{ADA}^{pred}$ | $\Delta E_{ADA} \pm \delta \Delta E_{ADA}$ |
| NDA | 800/880 | $0.539 \pm 0.022$ | — | — |
| | 800/960 | $0.532 \pm 0.020$ | — | — |
| | 880/960 | $0.530 \pm 0.023$ | — | — |
| ADN | 800/880 | $0.606 \pm 0.021$ | — | — |
| | 800/960 | $0.594 \pm 0.019$ | — | — |
| | 880/960 | $0.592 \pm 0.017$ | — | — |
| ADA | 800/880 | $0.744 \pm 0.014$ | $0.730 \pm 0.017$ | $0.014 \pm 0.031$ (0.022) |
| | 800/960 | $0.733 \pm 0.014$ | $0.722 \pm 0.016$ | $0.011 \pm 0.030$ (0.021) |
| | 880/960 | $0.739 \pm 0.019$ | $0.721 \pm 0.016$ | $0.018 \pm 0.035$ (0.025) |

We also computed the average FRET efficiency for each of the three constructs using Eq. 2, which incorporates measurements from a second excitation wavelength in the calculation. To assess the effects that different excitation wavelengths have on photobleaching, we performed separate experiments using three pairs of excitation wavelengths: 800 nm/880 nm, 800 nm/960 nm, and 880nm/960 nm. The results obtained from all three pairs of excitation wavelengths at the lowest excitation power (of 15 mW/pixel) are summarized in Table 2. In this case, the differences between the FRET efficiency values measured for the ADA fusion protein and the ones predicted based on measurements of the NDA and ADN constructs (1.4%, 1.1%, and 1.8%, for the 800 nm/880 nm, 800 nm/960 nm, and 880nm/960 nm wavelength pairs, respectively) were even smaller than those obtained for the single-wavelength excitation method and well within the calculated error for these quantities.

By comparing the results shown in Table 2 to those shown in Table 1, it becomes evident that the FRET efficiencies extracted from the single-wavelength excitation method were consistently higher (by 5% or more) than those obtained from the dual-wavelength excitation scheme, for all the constructs investigated. These differences originated from an overestimation of the 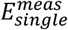, due to weak but non-negligible direct excitation of the acceptors at 800 nm, which is not accounted for in Eq. (1) but is included in Eq. (2) through the quantity *ρ*^*ex A*^ (see Supplementary Table 1 for values of this parameter).

Overall, the above observations support the notion that the use of the second excitation wavelength is warranted and confirm the validity of the measurement method based on two excitation wavelengths. Additional support for the validity of this method comes from the observation that the ratio of acceptor concentration to the total molecule concentration (i.e., molar fraction of the acceptors) obtained for low average excitation power conditions (15 mW) is about 0.64 ± 0.01, which closely matches the expected one of 0.67 for the ADA construct; this will be further discussed in the following section.

### 3.2 Effect of higher excitation powers on the FRET efficiency

Having established the accuracy of the method in the absence of significant photobleaching of the donors or acceptors, we next wanted to use it to explore the effect on the FRET efficiency values of increased excitation power and, hence, the likelihood that the molecules are photobleached. We performed measurements using three additional excitation powers (42 mW/point, 52 mW/point, and 62 mW/point) for both the single and the dual-wavelength excitation schemes. The measured FRET efficiency values of the ADA construct for all excitation wavelength pairs, presented in Fig. 3, decreased significantly with increasing the excitation power, due to photobleaching of either the donor or the acceptor. (The specific degree of photobleaching of the donors and acceptors has been assessed using computer simulations and will be discussed in the next section.) By contrast, the gray solid circles representing FRET efficiencies for single-wavelength excitation at 800 nm only show no significant effect of the increased power on FRET. The results for the other two FRET constructs (ADN and NDA) (Supplementary Tables 2-4) showed similar behavior for all the excitation wavelengths used. These changes suggest that the acceptors were photobleached during the measurement process through direct excitation, energy transfer, or both.

**Figure 3.**
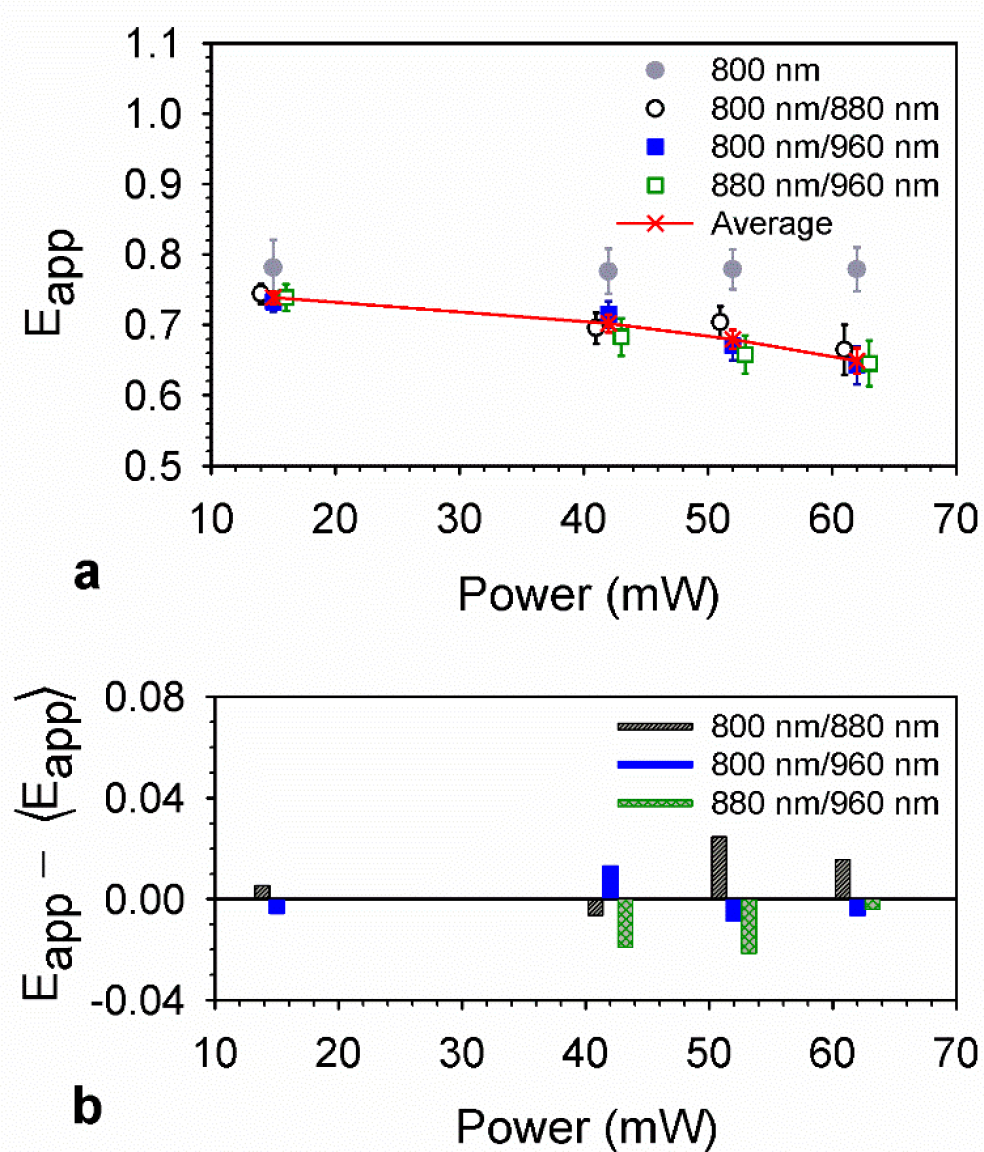
Apparent FRET efficiencies (*E*_*app*_) obtained using single- and dual-wavelength excitation methods at various powers for the cytoplasmic FRET construct ADA. Two excitation protocols were used: single-wavelength excitation at 800 nm, and dual-wavelength excitation at three different pairs of wavelengths (800 nm/880 nm, 800 nm/960 nm, and 880 nm/960 nm). **(a)** Measured *E*_*app*_ using various excitation schemes. The FRET efficiency for each excitation scheme was estimated from the weighted average FRET efficiencies of five different datasets obtained for that scheme at each power. **(b)** Difference between FRET efficiencies for each excitation wavelength pair and the weighted average FRET efficiencies across all excitation pairs indicated by the red dashed symbols in a. To enhance the legibility of the results, symbols for some measurement schemes were slightly shifted to the right or to the left of the actual power marked by the red symbol ×.

Interestingly, our analysis revealed (see Supplementary Tables 2-4) virtually no effect of the excitation power on the discrepancy between the measured (i.e., 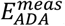) and predicted 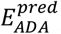 FRET efficiency for both excitation methods. In addition, for the dual-wavelength excitation method, the rate of FRET efficiency decrease with the increase in excitation power was similar for both the dimeric (NDA and ADN) and the trimeric (ADA) constructs (see Table 2 and Supplementary Tables 2-4). This is because, although currently the kinetic theory of FRET does not incorporate photobleaching effects, the contribution of photobleaching to dimeric FRET efficiencies is simply transferred to the FRET efficiency of the trimeric construct via the mathematical formula used (see Supplementary section SM2.1 and Ref [22]).

Also remarkably, for the single-wavelength excitation method using 800 nm laser light, the FRET efficiency remained the same for both the dimeric and the trimeric constructs across all four powers, although we have established that photobleaching must have occurred. By contrast, when the single excitation wavelength was 880 nm, the single-wavelength excitation method produced FRET efficiencies that increased with the average excitation power (see Supplementary Table 4). We hypothesize that these changes were caused by generation of acceptor-only complexes via photobleaching of donors, which were directly excited to a modest but significant degree (see above) by laser light at 800 nm. The additional signal produced by those acceptors contributed to the emission caused by excitation via energy transfer from the donors, thereby inadvertently compensating for the loss of signal due to photobleaching of acceptors residing within complexes with fluorescently active donors.

This hypothesis was easily tested and confirmed by computing the average molar fraction of the acceptors in the ADA complex, X_A_=[A]/{[A] +[D]}, from the same data used to determine the FRET efficiency (see Supplementary Method section SM1) for each excitation power. Indeed, while the observed X_A_ value for the construct ADA was 0.64 ± 0.01 in the absence of photobleaching at low excitation power (i.e., 15 mW/pixel), which closely approximates the expectation value of 0.67 for that construct, a significant increase in the observed XA value occurred for higher powers (see Fig. 4). That increase is consistent with an increase in the proportion of fluorescent protein constructs comprising only acceptors, which therefore qualitatively supports the observed constancy of (or even increase in) the FRET efficiency at different excitation powers as determined via the single-excitation wavelength method.

**Figure 4.**
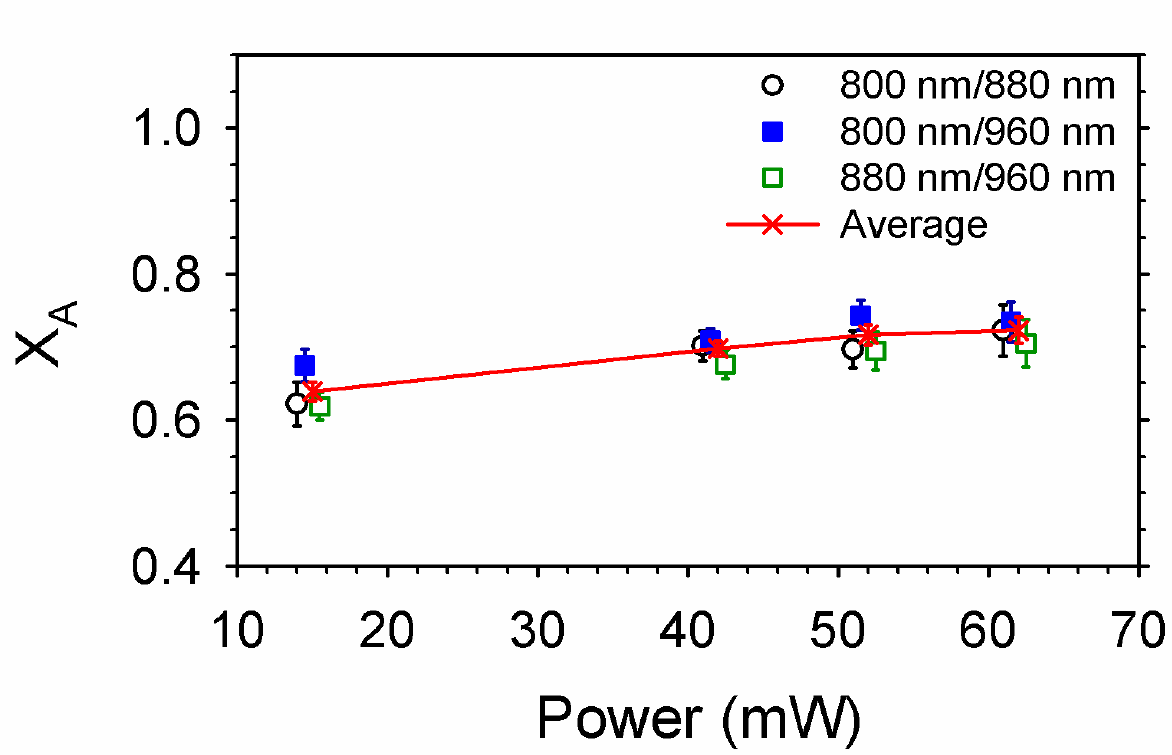
Acceptor molar fraction, X_A_=[A]/{[A] +[D]}, for a cytoplasmic FRET construct, ADA, transiently expressed in CHO cells, measured using different pairs of excitation wavelengths at different excitation powers. Black empty circles represent the X_A_ values estimated using the excitation pair 800 nm/880 nm. Blue solid squares represent the X_A_ values estimated using excitation pair 800 nm/960 nm, and green empty squares represent X_A_ values estimated using excitation pair 880 nm/960 nm. The red symbols × with red line represents the weighted average FRET efficiencies over all excitation wavelength pairs, i.e., the weighted average FRET efficiencies all the measurement performed with dual-wavelength excitation method. To enhance the legibility of the results, symbols for some excitation schemes were slightly adjusted shifted to the left and the right the power point marked by red symbol ×.

We will provide a more quantitative analysis of all the above observations in the next section.

### 3.3 Modeling the power dependence of the FRET efficiency using numerical simulations

To model the dependence of the measured FRET efficiency on the average excitation power (see Fig. 3), we performed numerical simulations to compute pixel-level FRET efficiencies for a population of FRET constructs residing at each pixel, which used a range of different photobleaching probabilities. The photobleaching probabilities represent the likelihood of a single fluorophore in the simulation transitioning from an active to a dark state for each excitation wavelength. The resultant FRET efficiency of the population of constructs residing at a pixel was recorded for each different set of photobleaching probabilities and compared to the actual measured FRET efficiency for the construct, using a fitting residual defined as the sum of the squares of the differences between measured and simulated FRET efficiencies (Equation SM16 in Supplemental Section SM4).

We generated three-dimensional plots with the three photobleaching probabilities (corresponding to D photobleaching during the first scan, A photobleaching during the first scan, and A photobleaching during the second scan) on the three axes and the fitting residual represented via a color code as the fourth dimension. Individual plots are displayed in Fig. 5 for three different sets of experimental powers (42 mW, 52 mW, and 62 mW) obtained for dual excitation at 800 nm and 960 nm (additional fitting residual distributinos generated for 800/880 nm and 880/960 nm wavelength pairs are displayed in Supplementary Figures SR1 and SR2). The lowest residual values correspond to the dark blue or black regions of the plots (see scale bar inset of Fig. 5). Fig. 6 compares the best-fit simulated FRET efficiencies, which incorporate the possibility of both D and A undergoing photobleaching at different probabilities, with the experimentally obtained values for each excitation power using the excitation wavelength pair of 800nm and 960nm (equivalent plots generated for 800/880 nm and 880/960 nm wavelength pairs are displayed in Supplementary Figures SR3 and SR4). The results showed a good agreement between simulations and experiments, indicating that the change in FRET efficiencies is explained well by differing photobleaching levels of D and A, which are collected in Table 3.

**Table 3.**
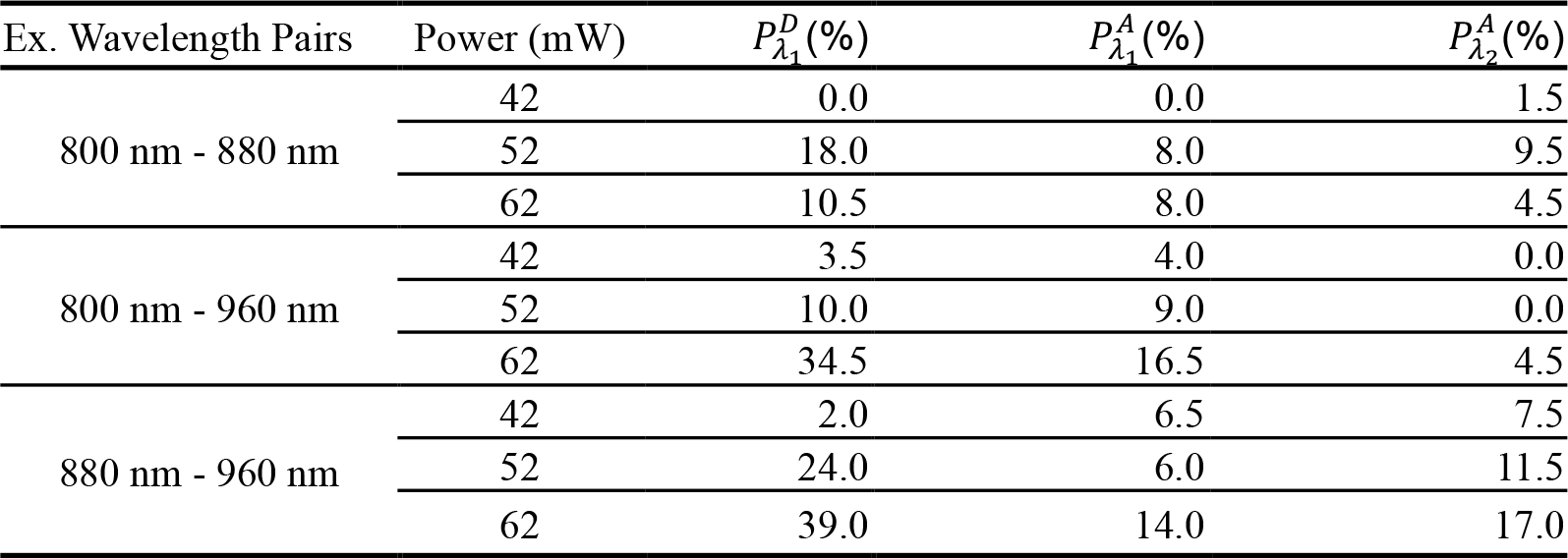
Summary of photobleaching probabilities, based on numerical simulation for three schemes of excitation pairs, at various excitation powers. The probabilities were estimated by fitting experimental results with simulated FRET efficiencies.

**Figure 5.**
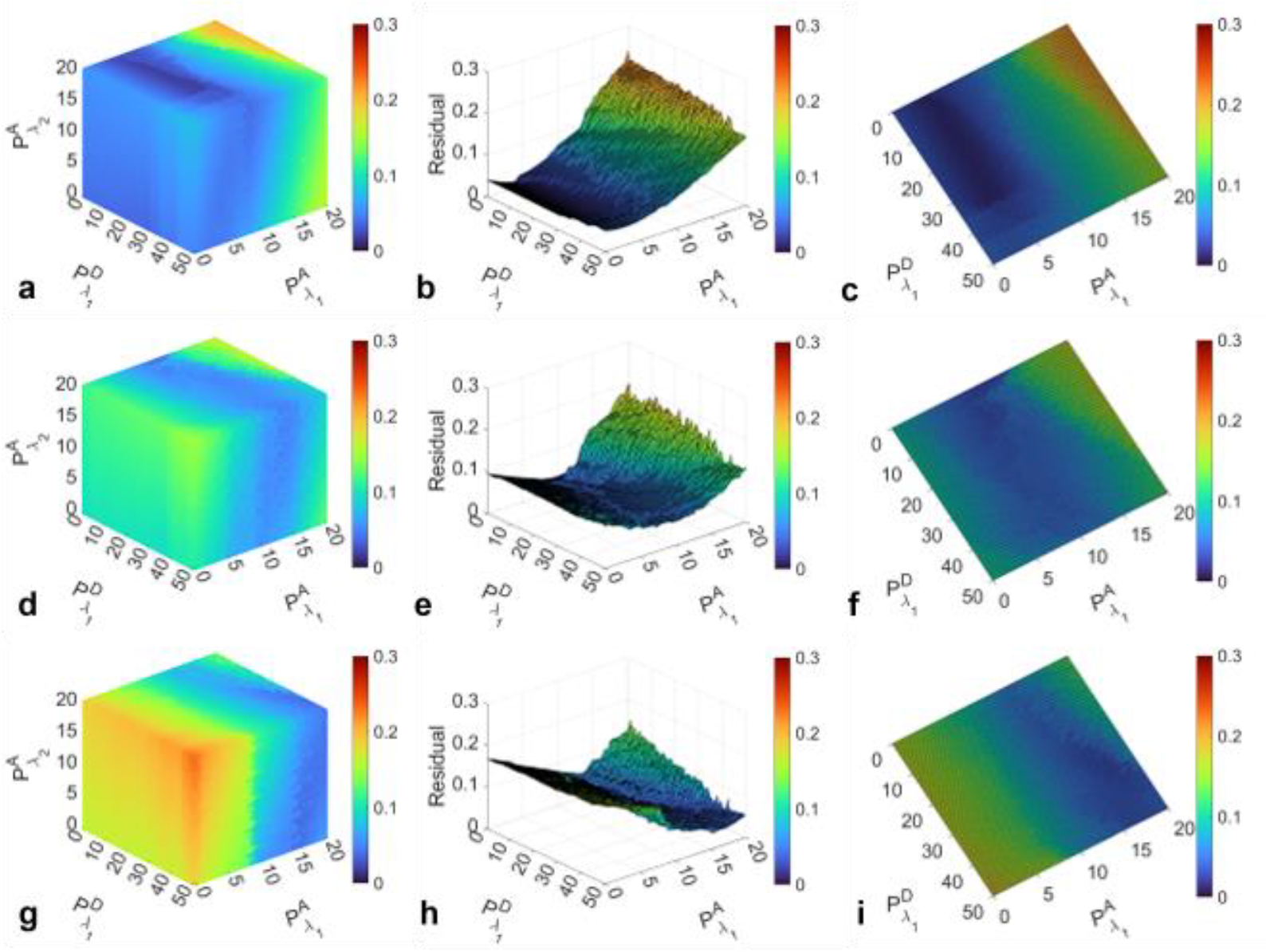
Systematic analysis of the agreement between simulated and measured FRET efficiencies (expressed as a fitting residual; see text) for a range of D and A photobleaching probabilities at 800 nm (*λ*_*1*_) and 960 nm (*λ*_*2*_) for three different excitation powers (increasing from top to bottom). (a, d&g) Pseudo-4D maps in which the magnitude of residuals, represented as different colors, is plotted against three different probabilities. **(b)** Cross-section of the pseudo-4D maps in a, showing the residual plot obtained by keeping the probability of acceptor bleaching at ***λ***_***2***_ fixed at a value corresponding to its minimum residual. **(c)** Top view of panel b, showing the residual distribution in a 2D map. The entire figure is organized into rows corresponding to different excitation powers: the top row **(panels a-c)** 42 mW/pixel; middle row **(panels d-f)** 52 mW/pixel; bottom row **(panels g-i)** 62 mW/pixel.

**Figure 6.**
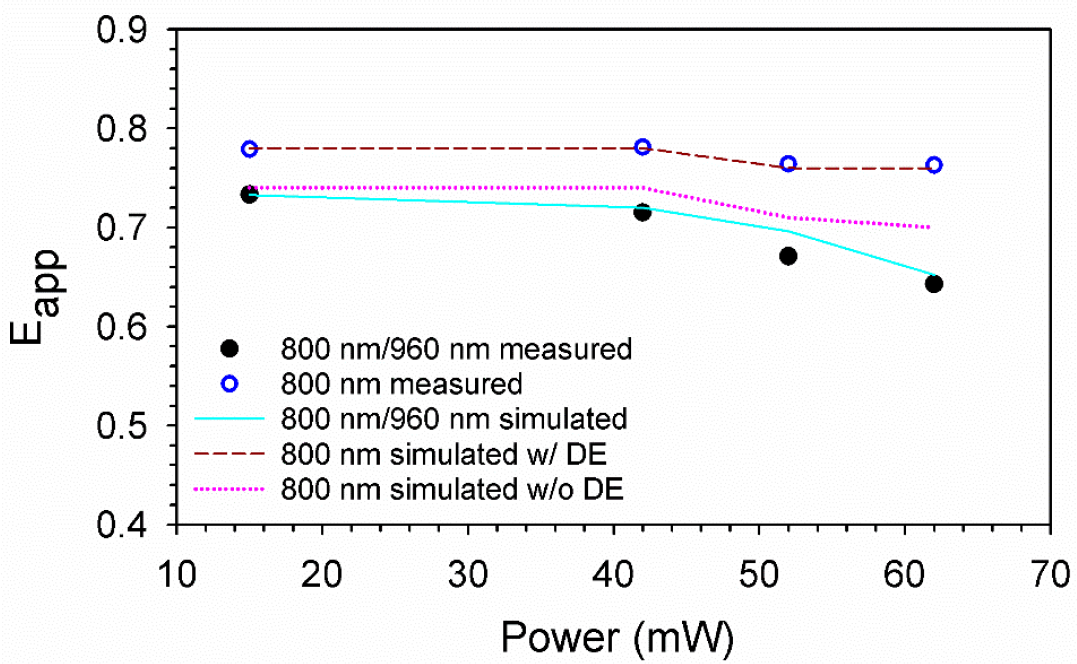
Comparison of measured and simulated FRET efficiencies (*E*_*app*_) for the cytoplasmic FRET construct, ADA, obtained using 800 nm/960 nm excitation at various powers. The black solid circles represent the measured FRET efficiencies using dual-wavelength excitation method with 800 nm/960 nm. Blue empty circles represent the measured FRET efficiencies for single-wavelength excitation at 800 nm. Whereas, cyan solid line represents the simulated FRET efficiencies curve for excitation pair 800 nm/960 nm. Dark red dashed line represents the simulated FRET efficiencies using single excitation at 800 nm with (w/) direct excitation (DE) of acceptor. Pink dotted line represents the simulated FRET efficiencies using single excitation at 800 nm without (w/o) direct excitation of acceptor.

Inspection of the photobleaching values in Table 3 indicates wide variability between different and even the same experimental conditions (such as the same *λ*_2_), due to the fact that the donor photobleaching at the second wavelength only affected an already small correction term in Eq. (2). Therefore, this minor effect is not discussed any further.

Overall, our simulations revealed a clear pattern of increasing photobleaching probability of both donor during the first scan at higher power, with donors exhibiting a wider range of photobleaching probabilities depending on the excitation power (see Table 3).

This analysis explained a second key trend observed in our experimental data, that the FRET efficiency values obtained from the single-wavelength excitation method remained constant regardless of excitation power. While this trend might initially suggest that the single-wavelength FRET values are impervious to photobleaching, a closer examination of the data shown in Fig. 6 reveals otherwise: Fig. 6 also displays the best-fit simulated FRET efficiency values obtained using the single-wavelength approach, with the contribution from acceptor direct excitation (DE) subtracted and plotted as a pink dotted line. This curve does indicate a monotonous decrease in *E*_*app*_ with increase in excitation power, which parallels the results of the dual-excitation scheme (compare the pink dotted line to the solid cyan line).

The apparent stability of the FRET efficiency across different excitation powers in the single-wavelength excitation method (brown dashed line) is deceiving and can be attributed to a balancing act: Increased excitation power leads to more donor photobleaching, which increases the number of free acceptors by inactivating the donor from the construct. This raises the measured FRET efficiency value due to the direct excitation of free acceptors. However, increased power also causes more acceptor photobleaching due to FRET, which tends to reduce FRET efficiency. Thus, in the single-wavelength excitation method, the increase in *E*_*app*_ caused by direct excitation balances out the decrease from acceptor photobleaching, resulting in a stable FRET efficiency across different power levels.

In contrast, in the dual-excitation method, corrections were made for the possible overestimation of FRET value caused by the direct excitation of the acceptor. However, acceptor photobleaching at the first excitation wavelength still leads to reduced FRET efficiency as excitation power increases. This is particularly relevant in FRET experiments, as FRET provides an extra pathway for acceptor photobleaching [37, 38].

## 4. CONCLUSION

One of the key findings in this study was the impact of the excitation power on FRET efficiency. With increasing the power, the FRET efficiency steadily decreased across all constructs and when measured and analyzed appropriately. This observation underlined the role of photobleaching in FRET experiments. Our constructs were comprised of the fluorescent proteins Cerulean (donor) and Venus (acceptor). Based on available information, the fluorescent protein Cerulean may be rather easily photobleached under direct excitation by light [25]. At the same time, while Venus is quite stable under direct excitation by light [25], in our study it was photobleached significantly through excitation by FRET, which is in agreement with the existing literature [37, 38].

The inadvertent decrease in FRET efficiency following excitation with relatively high laser powers (in our case, powers above 42 mW/pixel) may be erroneously interpreted as a larger distance between donor and acceptor molecules. This may have significant implications for applications where the actual distance between molecules is of interest, such as in studies focused on determination of the quaternary structure of protein oligomers [13, 14, 39]. Fortunately, at comparatively lower excitation powers, such as those corresponding to the first two datapoints in Fig. 6, both the single- and the dual-excitation wavelength methods provide results that are accurate within a few percent. To be meticulous, however, the single-wavelength excitation scheme may suffer from small systematic errors caused by the direct excitation of acceptors, which may be reduced by proper choice of acceptors.

Besides using low excitation powers, photobleaching may be reduced by reducing the dwell time of the excitation beam and developing fluorescent tags with better photostability. In addition, the present study also highlighted the importance of selecting the right excitation wavelengths. Excitation at 800 nm minimized direct excitation of the acceptor (Venus) while still efficiently exciting the donor (Cerulean). In contrast, the simulations indicated that the photobleaching of the acceptor was relatively high when using 880 nm as the first excitation wavelength compared to 800 nm. Furthermore, while results from the dual-wavelength excitation method confirmed a consistent trend of decreasing FRET efficiency with higher excitation power across all excitation wavelength pairs, the pair 800 nm/960 nm showed the most consistent overall weighted average FRET efficiency.

Another path forward would be to extend the kinetic theory of FRET, by incorporating photobleaching rates in addition to spontaneous emission and energy transfer rates, so that one can tease apart the contributions of these effects to the FRET efficiencies determined from data affected by photobleaching. Once available, this could be used together with methods described above for reducing photobleaching or just by itself. Such a theoretical undertaking will require a good understanding of all the photophysical and photochemical phenomena involved. This process, initiated through the computer simulations presented in this paper, needs more rigorous treatment in the future and will benefit from analysis of existing [37, 38] and future experimental studies. This will allow experimentalists to use the theory as a means of obtaining correct results from experiments rather than as a toll of assessing the errors introduced by harsh experimental conditions.

## Supporting information

Supplementary Methods and Results

## Notes

### Competing Interest Statement

The authors have declared no competing interest.

