## Supplementary Methods and Results for "Impact of photobleaching of fluorescent proteins on FRET measurements under two-photon excitation"

Dhruba P. Adhikari[1], Michael R. Stoneman[1], Gabriel Biener[1], and Valerica Raicu[1]

[1]*Department of Physics, , University of Wisconsin-Milwaukee, Milwaukee, WI, USA*

### SUPPLEMENTARY METHODS

#### SM1. Determination of the acceptor mole fraction ($X_A$)

Four quantities were extracted from the micro-spectroscopic scans of a sample at two excitation wavelengths: the fluorescence of the donor and acceptor in the presence of each other at the first excitation wavelength ($F^{DA}(\lambda_{ex,2})$ and $F^{AD}(\lambda_{ex,2})$, respectively) as well as at the second excitation wavelength ($F^{DA}(\lambda_{ex,2})$ and $F^{AD}(\lambda_{ex,2})$). From these measured quantities, we determined the total donor emission at the first excitation wavelength as if there were no FRET present [2, 3]:

$$F^D(\lambda_{ex,1}) = F^{DA}(\lambda_{ex,1}) + \frac{Q^D}{Q^A} \cdot F^{AD}(\lambda_{ex,1}) - \rho^A \frac{Q^D}{Q^A} \cdot F^A(\lambda_{ex,2}), \quad \text{(SM1)}$$

and the total acceptor fluorescence at the second excitation wavelength in the absence of FRET:

$$F^A(\lambda_{ex,2}) = \left[\frac{\rho^D \cdot F^{AD}(\lambda_{ex,2}) - F^{AD}(\lambda_{ex,1})}{\rho^D - \rho^A}\right] \quad \text{(SM2)}$$

The intensity ratios of the donors and acceptors $\rho^{ex,\,D}$ and $\rho^{ex,\,A}$, respectively, between the two excitation wavelengths, (i.e., $\rho^{ex,y} = F_1^y / F_2^y$ where *y=D or A)* were obtained by scanning samples expressing only the donor or only the acceptor using the first and second excitation wavelengths.

The average values of $F^D(\lambda_{ex,1})$ and $F^A(\lambda_{ex,2})$ over a given segment were used to estimate the acceptor mole fraction ($X_A$) for the given segment. $X_A$ was calculated based on the number of donors, $n_D$, and acceptors, $n_A$, in the segment, using the following equation:

$$X_A = \frac{n_D}{n_n + n_D}, \quad \text{(SM3)}$$

The xxpressions for the for the total number of donors and acceptors residing in a given pixel, $n_D$ and $n_A$, respectively, were determined as follows [2, 3]:

$$n_D = \frac{F^D(\lambda_{ex,1}) \cdot V_{TPE}(\lambda_{ex,1}) \cdot N_A}{\sigma_{2P}^D(\lambda_{ex,1}) \cdot Q^D \cdot CE}, \quad \text{(SM4)}$$

$$n_A = \frac{F^A(\lambda_{ex,2}) \cdot V_{TPE}(\lambda_{ex,2}) \cdot N_A}{\sigma_{2P}^A(\lambda_{ex,2}) \cdot Q^A \cdot CE}, \quad \text{(SM5)}$$

where $Q^D$ and $Q^A$ are the quantum yields of the donor and acceptor, respectively. $N_A$ represents Avogadro's number, $V_{TPE}(\lambda_{ex})$ represents the two-photon excitation volume [4, 5] at a specific excitation wavelength, $\lambda_{ex}$, and $CE$ the instrument specific photon collection efficiency. Additionally, $\sigma_{2P}^{D}(\lambda_{ex,1})$ represents the 2P absorption cross section of the donor at the first excitation wavelength and $\sigma_{2P}^{A}(\lambda_{ex,2})$ the 2P absorption cross section of the acceptor at the second wavelength. Inserting equations SM12 and SM13 into SM11 gives:

$$X_A = \frac{F^A(\lambda_{ex,2})}{F^A(\lambda_{ex,2}) + F^D(\lambda_{ex,1})\dfrac{\sigma_{2P}^{A}(\lambda_{ex,2}) \cdot Q^A}{\sigma_{2P}^{D}(\lambda_{ex,1}) \cdot Q^D}}, \qquad \text{(SM6)}$$

For our analysis, we adjusted peak 2P absorption cross-sections from previous reports based on the measured fraction of absorption at a specific excitation wavelength compared to the peak absorption. Specifically, we used a value of 13 GM at 858 nm for Ceurlean [6] and 19.0 GM at 965 nm for Venus [7]. The absorption fraction at each excitation wavelength, relative to the maximum absorption wavelength, was determined from excitation spectra obtained from cells expressing only Cerulean or only Venus using our instrument.

### SM2. Derivation of the relationship between FRET dimers and trimers

#### *SM2.1. Summary of the kinetic theory of FRET*

The FRET efficiency of each individual donor, *i*, in a protein complex containing *n* protomers, with *k* donors and *n-k* acceptors, is a function of the energy transfer rates occurring between the donor and each of the acceptors in the complex, according to the following expression [1]:

$$E_{i,k,n} = \frac{\sum_{j=1}^{n-k} \Gamma_{i,j}^{FRET}/(\Gamma^{r,D} + \Gamma^{nr,D})}{1 + \sum_{j=1}^{n-k} \Gamma_{i,j}^{FRET}/(\Gamma^{r,D} + \Gamma^{nr,D})}, \qquad \text{(SM7)}$$

where the summation index *j* represents a sum over all of the acceptors present in the oligomeric complex. $\Gamma^{r,D}$ and $\Gamma^{nr,D}$ are the rate constants of de-excitation through radiative (i.e., photon emission) and non-radiative (e.g., internal conversion) processes, respectively, and $\Gamma_{i,j}^{FRET}$ is the rate constant of de-excitation through FRET between the $i^{th}$ donor and $j^{th}$ acceptor. The apparent FRET efficiency for a given complex can be predicted by averaging the $E_{i,k,n}$ values from each individual donor:

$$E^{pred} = \frac{1}{k}\sum_{i=1}^{k} E_{i,k,n} = \frac{1}{k}\sum_{i=1}^{k} \frac{\sum_{j=1}^{n-k} \Gamma_{i,j}^{FRET}/(\Gamma^{r,D} + \Gamma^{nr,D})}{1 + \sum_{j=1}^{n-k} \Gamma_{i,j}^{FRET}/(\Gamma^{r,D} + \Gamma^{nr,D})}. \qquad \text{(SM8)}$$

If the protein complex contains only one donor ($k = 1$), as is the case for all of the constructs in this study, Eq.SM2 simplifies to[1]:

$$E^{pred} = \frac{\sum_{j=1}^{n-1} \Gamma_{j}^{FRET}/(\Gamma^{r,D} + \Gamma^{nr,D})}{1 + \sum_{j=1}^{n-1} \Gamma_{j}^{FRET}/(\Gamma^{r,D} + \Gamma^{nr,D})}. \qquad \text{(SM9)}$$

#### *SM2.2. Computing $E_{app}$ for multimeric constructs from individual D-A energy transfer rates*

For multiplexes containing more than one acceptor, such as the ADA complex measured in this study, the predicted FRET efficiency of Eq. SM3 can be calculated if all the individual rates of energy transfer between the donor and each acceptor, $\Gamma_j^{FRET}$, are known. The two individual rates of energy transfer present in the ADA complex were found by measuring the FRET efficiency of the NDA and ADN constructs. The relationship between the FRET efficiency of a complex containing a single D and A and the rate of energy transfer between D and A is given as:

$$E_j = \frac{\Gamma_j^{FRET}}{\Gamma^{r,D} + \Gamma^{nr,D} + \Gamma_j^{FRET}}, \quad \text{(SM10)}$$

Combining Eq. SM3 with Eq. SM4 results in:

$$E^{pred} = \frac{\sum_{j=1}^{n-1} E_j/1 - E_j}{1 + \sum_{j=1}^{n-k} E_j/1 - E_j}, \quad \text{(SM11)}$$

The predicted FRET efficiency for the trimeric FRET construct, $E_{ADA}^{pred}$ , was determined by inserting the values of the measured FRET efficiency ($E_j$) of dimeric FRET constructs (i.e., NDA and ADN) into Eq. SM5, as follows:

$$E_{ADA}^{pred} = \frac{E_{NDA}^{meas}/(1 - E_{NDA}^{meas}) + E_{ADN}^{meas}/(1 - E_{ADN}^{meas})}{1 + E_{NDA}^{meas}/(1 - E_{NDA}^{meas}) + E_{ADN}^{meas}/(1 - E_{ADN}^{meas})}, \quad \text{(SM12)}$$

where $E_{NDA}^{meas}$ and $E_{ADN}^{meas}$ represent the measured FRET efficiencies of dimeric FRET constructs NDA and ADN, respectively.

The uncertainty associated with each of the measured FRET efficiency values was taken to be one standard deviation of the distribution of FRET values obtained from all segments of the associated sample. The error associated with the predicted FRET value of the ADA complex, $\delta E_{ADA}^{pred}$, was found using linear error propagation of the individual NDA and ADN measurements as follows:

$$\delta E_{ADA}^{pred} = \frac{\frac{\sigma_{E_{NDA}}}{(1 - E_{NDA}^{meas})^2} + \frac{\sigma_{E_{ADN}}}{(1 - E_{ADN}^{meas})^2}}{\left(\frac{E_{NDA}^{meas}}{1 - E_{NDA}^{meas}} + \frac{E_{ADN}^{meas}}{1 - E_{ADN}^{meas}}\right)\left(1 + \frac{E_{NDA}^{meas}}{(1 - E_{NDA}^{meas})} + \frac{E_{ADN}^{meas}}{(1 - E_{ADN}^{meas})}\right)}, \quad \text{(SM13)}$$

where $\sigma_{E_{NDA}}$ and $\sigma_{E_{ADN}}$ are the standard deviation (SD) of the measured FRET efficiency distributions obtained for the constructs NDA and ADN, respectively.

A value of $E_{ADA}^{pred}$ was calculated for each excitation condition (i.e., for various excitation wavelengths and powers) using the values obtained from separate measurements on the constructs NDA and ADN using the exact same imaging conditions.

**SM3. Calculation of the weighted average and standard error of measured $E_{app}$**

Four to five datasets were collected for each of the three FRET constructs at each of the excitation powers and pairs of excitation wavelengths. Those results were used to compute a weighted average for each construct for a specific power and excitation wavelength pair using following equation:

$$E = \frac{\Sigma_m \frac{1}{\sigma_m^2} E_m}{\Sigma_m \frac{1}{\sigma_m^2}}, \qquad \text{(SM14)}$$

where, $E_m$ and $\sigma_m$ represent the average and standard deviation, respectively, of the FRET efficiency distribution for each dataset, m.

Similarly, the standard error associated with each of the weighted FRET efficiency values was obtained from the weighted standard deviation of the corresponding FRET values, using following equation:

$$\mathrm{SD_w} = \left(\Sigma_m \frac{1}{\sigma_m^2}\right)^{-\frac{1}{2}}. \qquad \text{(SM15)}$$

### SM4. Flowcharts illustrating workflow of the data-fitting procedure

An iterative data-fitting procedure was implemented to determine the photobleaching probabilities of the donors and acceptors at each excitation wavelength and power. The procedure iteratively applied an array of photobleaching parameters to the input of a numerical simulation of FRET complexes (workflow of the numerical simulation illustrated in Supplemental Fig. SM1). For each set of photobleaching parameters used as input, a FRET efficiency value was determined from the simulation. These numerically simulated FRET efficiency values, incorporating photobleaching probabilities, were compared with the experimentally measured FRET efficiencies by computing the fitting residual

$$Residual = \left[\sum_i \left(E^i_{mea,single} - E^i_{sim,single}\right)^2 + \sum_i \left(E^i_{mea,dual} - E^i_{sim,dual}\right)^2\right]^2, \quad \text{(SM16)}$$

where $i$ is a summation index that stands for the constructs "NDA," "ADN," and "ADA," $E^i_{mea,single}$ and $E^i_{sin,single}$ are FRET efficiencies measured using the single-wavelength protocol and simulated, respectively, for each construct, while $E^i_{m,dual}$ and $E^i_{s,dual}$ are FRET efficiencies measured using the dual-wavelength excitation protocol and, respectively, simulated for each construct. The input photobleaching parameters were varied from 0 to 1 in steps of 0.5, as illustrated in Supplementary Fig. SM2. The photobleaching probabilities of D and A for a particular excitation power and excitation wavelength pair were determined based on the set of photobleaching probabilities corresponding to the lowest overall residual.

**$F^{DA}$ and $F^{AD}$ calculation based on input values**

| Construct | **ADA** | **ADN** | **NDA** | NDN | ANA | ANN | NNA | NNN |
|---|---|---|---|---|---|---|---|---|
| $E_{app}$ | 0.739 | 0.597 | 0.534 | 0.000 | 0.000 | 0.000 | 0.000 | 0.000 |
| $F_1^{DA}$ | 0.144 | 0.223 | 0.257 | 0.552 | 0.000 | 0.000 | 0.000 | 0.000 |
| $F_1^{AD}$ | 0.473 | 0.352 | 0.320 | 0.000 | 0.098 | 0.049 | 0.049 | 0.000 |
| $F_2^{AD}$ | 1.519 | 0.969 | 0.911 | 0.000 | 0.840 | 0.420 | 0.420 | 0.000 |

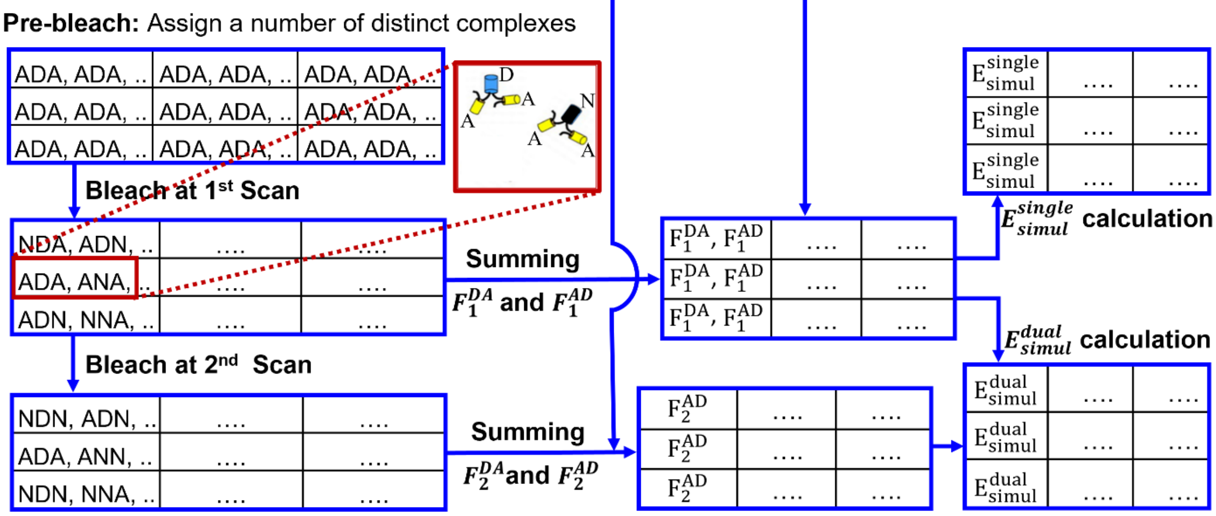


**Supplemental Figure SM1. Flowchart illustrating the step-by-step process of the numerical simulation for analyzing the effect of photobleaching on the FRET efficiency of a trimeric FRET construct, ADA.** A step-by-step description of the simulation is given in Section 2.5 of the manuscript.

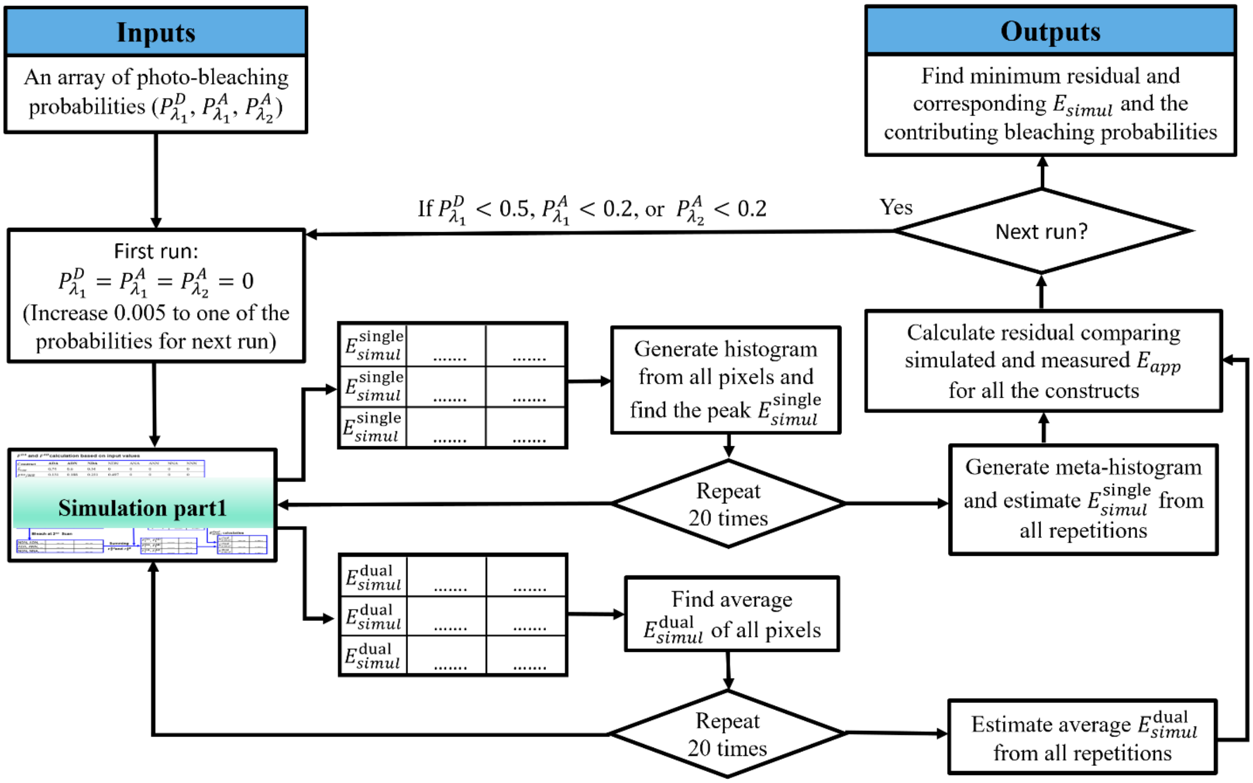


**Supplemental Figure SM2. Flowchart illustrating the data fitting procedure for determining donor and acceptor photobleaching probabilities under a specific excitation condition.** The procedure applied an array of photobleaching parameters to a numerical simulation of FRET complexes (workflow for the numerical simulation shown in Supplemental Fig. SM2). For each set of parameters, a FRET efficiency value was determined. These values, incorporating photobleaching probabilities, were compared with experimental FRET efficiencies using the fitting residual (Supplemental Eq. SM16). The set of parameters resulting in the minimum residual were taken as the specific contributions of donor and acceptor photobleaching in the FRET measurement under the excitation condition.

### SUPPLEMENTARY RESULTS

**Supplemental Table 1. The intensity ratios of the donors ($\rho^{ex,D}$) and acceptors ($\rho^{ex,A}$) between two excitation wavelengths at various excitation powers.** The measurements were performed for three different excitation wavelength pairs: 800 nm/880 nm, 800 nm/960 nm, and 880 nm/960 nm and four different excitation powers: 15 mW/point, 42 mW/point, 52 mW/point, and 62 mW/point.

| Ex. Wavelengths | Cofficient | 15 mW/point | 42 mW/point | 52 mW/point | 62 mW/point |
|---|---|---|---|---|---|
| 800 nm /880 nm | $\rho^{ex,D}$ | 0.46 | 0.55 | 0.60 | 0.61 |
| | $\rho^{ex,A}$ | 0.1 | 0.14 | 0.15 | 0.20 |
| 800 nm /960 nm | $\rho^{ex,D}$ | 11.45 | 8.33 | 8.37 | 7.17 |
| | $\rho^{ex,A}$ | 0.05 | 0.05 | 0.07 | 0.08 |
| 880 nm /960 nm | $\rho^{ex,D}$ | 23.91 | 19.06 | 17.14 | 13.73 |
| | $\rho^{ex,A}$ | 0.37 | 0.42 | 0.43 | 0.46 |

**Supplemental Table 2. Weighted average of apparent FRET efficiencies ± SE of all datasets obtained for each FRET construct measured using both single and dual excitation methods for the experiments carried out at 800 and 880 nm.** The following average excitation powers per point were used: 15 mW/point, 42 mW/point, 52 mW/point, and 62 mW/point. See Supplemental equations SM6-SM8 for the formulas used to compute weighted averages and SE.

| Power (mW) | Constructs | Excitation Wavelength (nm) | Apparent FRET efficiencies | | |
|---|---|---|---|---|---|
| | | | $E^{meas} \pm SD_w$ | $E_{ADA}^{pred} \pm \delta E_{ADA}^{pred}$ | $\Delta E_{ADA} \pm \delta\Delta E_{ADA}$ |
| 15 | NDA | 800 | 0.556 ± 0.023 | — | — |
| | | 800/880 | 0.539 ± 0.022 | — | — |
| | ADN | 800 | 0.631 ± 0.021 | — | — |
| | | 800/880 | 0.606 ± 0.021 | — | — |
| | ADA | 800 | 0.778 ± 0.026 | 0.748 ± 0.017 | 0.030 ± 0.043 (0.031) |
| | | 800/880 | 0.744 ± 0.014 | 0.730 ± 0.017 | 0.014 ± 0.031 (0.022) |
| 42 | NDA | 800 | 0.568 ± 0.019 | — | — |
| | | 800/880 | 0.504 ± 0.020 | — | — |
| | ADN | 800 | 0.629 ± 0.020 | — | — |
| | | 800/880 | 0.571 ± 0.020 | — | — |
| | ADA | 800 | 0.773 ± 0.026 | 0.751 ± 0.015 | 0.022±0.041 (0.030) |
| | | 800/880 | 0.696 ± 0.022 | 0.701 ± 0.017 | -0.005±0.039 (0.028) |
| 52 | NDA | 800 | 0.549 ± 0.029 | — | — |
| | | 800/880 | 0.471 ± 0.027 | — | — |
| | ADN | 800 | 0.635 ± 0.020 | — | — |
| | | 800/880 | 0.550 ± 0.018 | — | — |
| | ADA | 800 | 0.786 ± 0.025 | 0.747 ± 0.019 | 0.039±0.044 (0.031) |
| | | 800/880 | 0.704 ± 0.022 | 0.679 ± 0.019 | 0.025 ± 0.041 (0.029) |
| 62 | NDA | 800 | 0.554 ± 0.026 | — | — |
| | | 800/880 | 0.452 ± 0.033 | — | — |
| | ADN | 800 | 0.637 ± 0.021 | — | — |
| | | 800/880 | 0.517 ± 0.024 | — | — |
| | ADA | 800 | 0.776 ± 0.028 | 0.750 ± 0.018 | 0.026 ± 0.046 (0.033) |
| | | 800/880 | 0.665 ± 0.035 | 0.655 ± 0.025 | 0.010 ± 0.060 (0.043) |

**Supplemental Table 3. Weighted average of apparent FRET efficiencies ± SE of all datasets obtained for each FRET construct measured using both single and dual excitation methods for the experiments carried out at 800 and 960 nm.** The following average excitation powers per point were used: 15 mW/point, 42 mW/point, 52 mW/point, and 62 mW/point. See Supplemental equations SM6-SM8 for the formulas used to compute weighted averages and SE.

| Power (mW) | Constructs | Excitation Wavelength (nm) | Apparent FRET efficiencies | | |
|---|---|---|---|---|---|
| | | | $E_{meas}$± $SD_w$ | $E_{pred}^{meas}$± $\delta E_{pred}$ | $\Delta E^{meas}$± $\delta\Delta E$ |
| 15 | NDA | 800 | 0.575 ± 0.021 | — | — |
| | | 800/960 | 0.532 ± 0.020 | — | — |
| | ADN | 800 | 0.615± 0.017 | — | — |
| | | 800/960 | 0.594 ± 0.019 | — | — |
| | ADA | 800 | 0.779 ± 0.048 | 0.747 ± 0.015 | 0.032 ± 0.063 (0.050) |
| | | 800/960 | 0.733 ± 0.014 | 0.722 ± 0.016 | 0.011 ± 0.030 (0.021) |
| 42 | NDA | 800 | 0.575 ± 0.020 | — | — |
| | | 800/960 | 0.503 ± 0.019 | — | — |
| | ADN | 800 | 0.629 ± 0.017 | — | — |
| | | 800/960 | 0.577 ± 0.016 | — | — |
| | ADA | 800 | 0.781 ± 0.035 | 0.753 ± 0.014 | 0.028 ± 0.049 (0.038) |
| | | 800/960 | 0.715 ± 0.018 | 0.704 ± 0.015 | 0.011 ± 0.033 (0.023) |
| 52 | NDA | 800 | 0.583 ± 0.026 | — | — |
| | | 800/960 | 0.470 ± 0.027 | — | — |
| | ADN | 800 | 0.646 ± 0.018 | — | — |
| | | 800/960 | 0.556 ± 0.021 | — | — |
| | ADA | 800 | 0.764 ± 0.023 | 0.763 ± 0.016 | 0.001 ± 0.039 (0.028) |
| | | 800/960 | 0.671 ± 0.021 | 0.681 ± 0.021 | -0.010 ± 0.042 (0.029) |
| 62 | NDA | 800 | 0.557 ± 0.024 | — | — |
| | | 800/960 | 0.428 ± 0.027 | — | — |
| | ADN | 800 | 0.623 ± 0.020 | — | — |
| | | 800/960 | 0.513 ± 0.024 | — | — |
| | ADA | 800 | 0.763 ± 0.030 | 0.744 ± 0.017 | 0.019 ± 0.047 (0.035) |
| | | 800/960 | 0.643 ± 0.035 | 0.643 ± 0.023 | -0.000 ± 0.058 (0.042) |

**Supplemental Table 4. Weighted average of apparent FRET efficiencies ± SE of all datasets obtained for each FRET construct measured using both single and dual excitation methods for the experiments carried out at 880 and 960 nm.** The following average excitation powers per point were used: 15 mW/point, 42 mW/point, 52 mW/point, and 62 mW/point. See Supplemental equations SM6-SM8 for the formulas used to compute weighted averages and SE.

| Power (mW) | Constructs | Excitation Wavelength (nm) | Apparent FRET efficiencies | | |
|---|---|---|---|---|---|
| | | | $E^{meas} \pm \mathrm{SD_w}$ | $E_{ADA}^{\mathrm{pred}} \pm \delta E_{ADA}^{pred}$ | $\Delta E_{ADA} \pm \delta\Delta E_{ADA}$ |
| 15 | NDA | 880 | 0.649 ± 0.019 | — | — |
| | | 880/960 | 0.530 ± 0.023 | — | — |
| | ADN | 880 | 0.713 ± 0.015 | — | — |
| | | 880/960 | 0.592 ± 0.017 | — | — |
| | ADA | 880 | 0.835 ± 0.032 | 0.812 ± 0.012 | 0.023 ± 0.044 (0.034) |
| | | 880/960 | 0.739 ± 0.019 | 0.721 ± 0.016 | 0.018 ± 0.035 (0.025) |
| 42 | NDA | 880 | 0.677 ± 0.017 | — | — |
| | | 880/960 | 0.477 ± 0.023 | — | — |
| | ADN | 880 | 0.713 ± 0.015 | — | — |
| | | 880/960 | 0.565 ± 0.020 | — | — |
| | ADA | 880 | 0.839 ± 0.019 | 0.821 ± 0.011 | 0.018 ± 0.030 (0.022) |
| | | 880/960 | 0.683 ± 0.026 | 0.688 ± 0.018 | -0.005 ± 0.044 (0.032) |
| 52 | NDA | 880 | 0.670 ± 0.020 | — | — |
| | | 880/960 | 0.460 ± 0.026 | — | — |
| | ADN | 880 | 0.732 ± 0.016 | — | — |
| | | 880/960 | 0.536 ± 0.021 | — | — |
| | ADA | 880 | 0.849 ± 0.019 | 0.826 ± 0.012 | 0.022 ± 0.031 (0.022) |
| | | 880/960 | 0.658 ± 0.027 | 0.667 ± 0.021 | -0.010 ± 0.048 (0.034) |
| 62 | NDA | 880 | 0.680 ± 0.019 | — | — |
| | | 880/960 | 0.436 ± 0.030 | — | — |
| | ADN | 880 | 0.716 ± 0.014 | — | — |
| | | 880/960 | 0.513 ± 0.024 | — | — |
| | ADA | 880 | 0.850 ± 0.020 | 0.823 ± 0.011 | 0.027 ± 0.031 (0.023) |
| | | 880/960 | 0.645 ± 0.033 | 0.646 ± 0.024 | -0.001 ± 0.057 (0.041) |

**Supplemental Table 5. The input FRET efficiencies for numerical simulation. These values are weighted average FRET efficiencies across all excitation pairs at power 15 mW/point.**

| Constructs | Avg. $E_{meas}^{dual}$ of all ex. pairs at 15 mW/point |
|---|---|
| NDA | 0.534 |
| NDA | 0.597 |
| ADA | 0.739 |

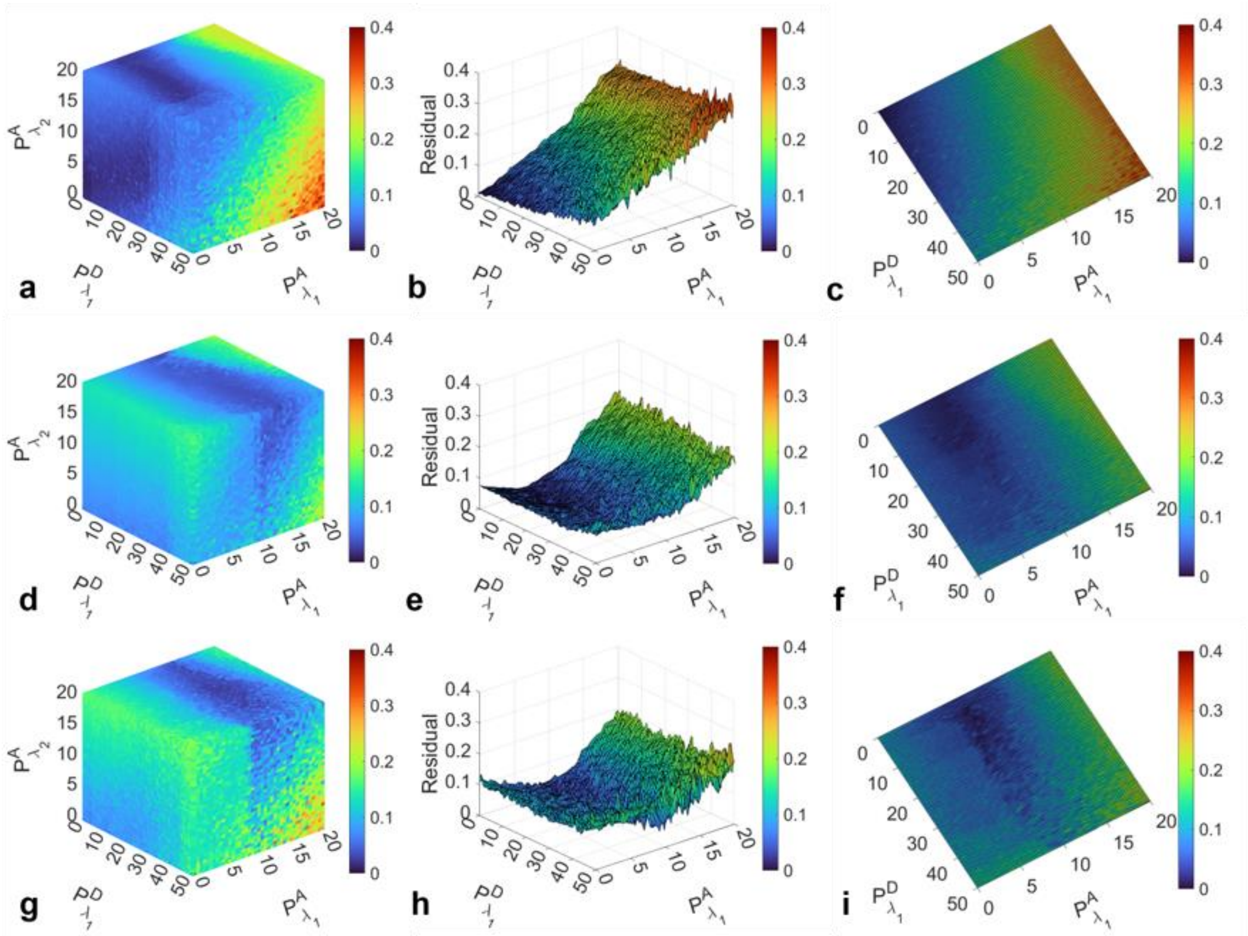


**Supplemental Figure SR1. Distribution of fitting residual as a function of photobleaching probabilities, calculated comparing simulated and measured FRET efficiencies of all three constructs under 800 nm/880 nm excitation for three different excitation powers. (a)** 3D residual map that show probability of donor bleaching at the first wavelength along x-axis, the probability of acceptor bleaching at the first wavelength along y-axis, and the probability of acceptor bleaching at a second wavelength along z-axis, with color gradients indicating the magnitude of residuals. **(b)** The modifies version of panel a by substituting the vertical axis with residual values, offering a refined analysis of residual distribution keeping the probability of acceptor bleaching at second wavelength fixed corresponding to minimum residual. **(c)** Top view of panel b, highlighting residual distribution in 2D map. The figure is organized into rows corresponding to different excitation powers: the top row (a-c) for excitation power 42 mW/point, the middle row (d-f) for 52 mW/point, and the bottom row (g-i) for 62 mW/point, showcasing how variations in excitation power influence the photobleaching dynamics and FRET efficiency residuals.

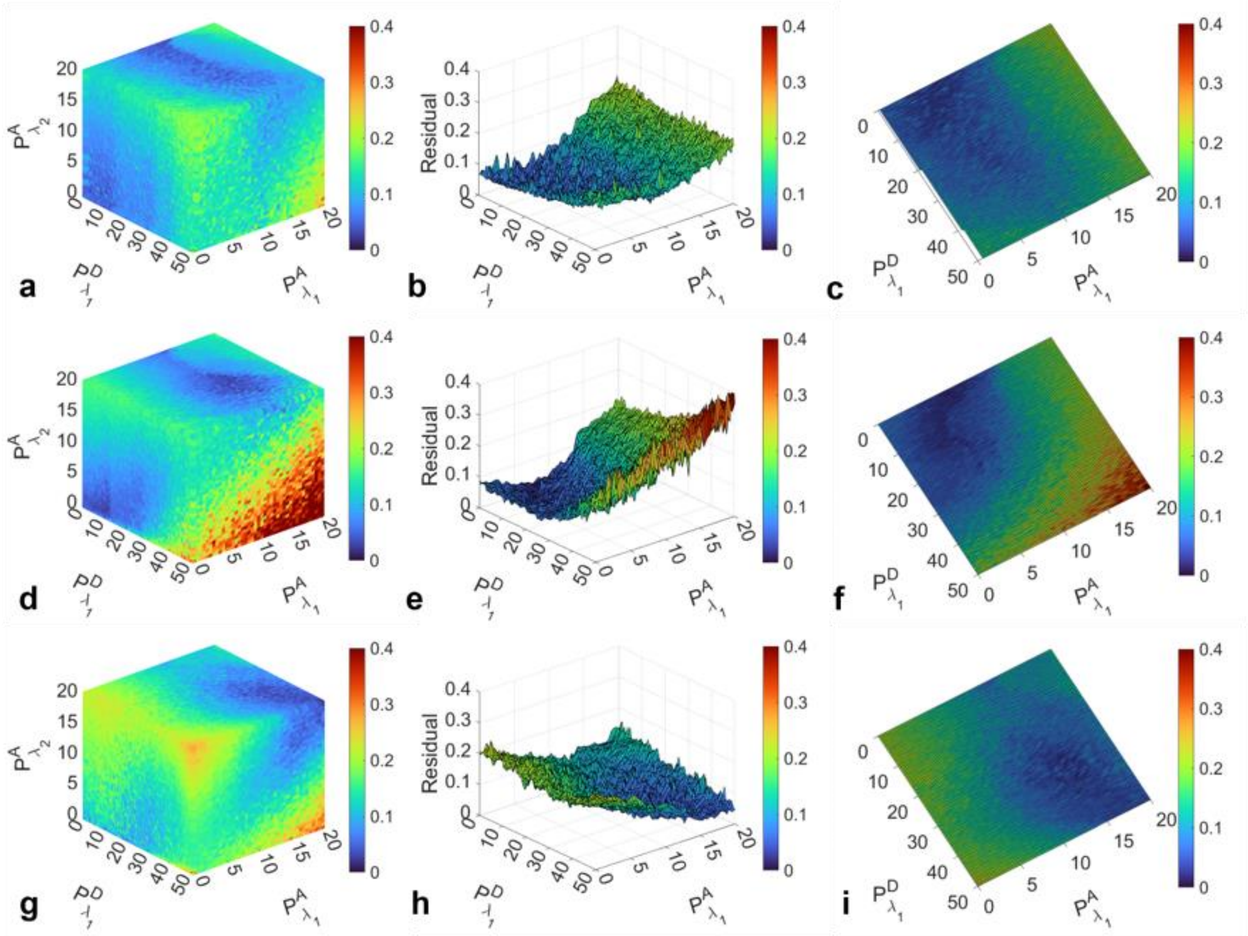


**Supplemental Figure SR2. Distribution of fitting residual as a function of photobleaching probabilities, calculated comparing simulated and measured FRET efficiencies of all three constructs under 880 nm/960 nm excitation for three different excitation powers. (a)** 3D residual map that show probability of donor bleaching at the first wavelength along x-axis, the probability of acceptor bleaching at the first wavelength along y-axis, and the probability of acceptor bleaching at a second wavelength along z-axis, with color gradients indicating the magnitude of residuals. **(b)** The modifies version of panel a by substituting the vertical axis with residual values, offering a refined analysis of variance between measured and simulated data keeping the probability of acceptor bleaching at second wavelength fixed corresponding to minimum residual. **(c)** Top view of panel b, highlighting residual distribution in 2D map. The figure is organized into rows corresponding to different excitation powers: the top row (a-c) for excitation power 42 mW/point, the middle row (d-f) for 52 mW/point, and the bottom row (g-i) for 62 mW/point, showcasing how variations in excitation powers influence the photobleaching dynamics and FRET efficiency residuals

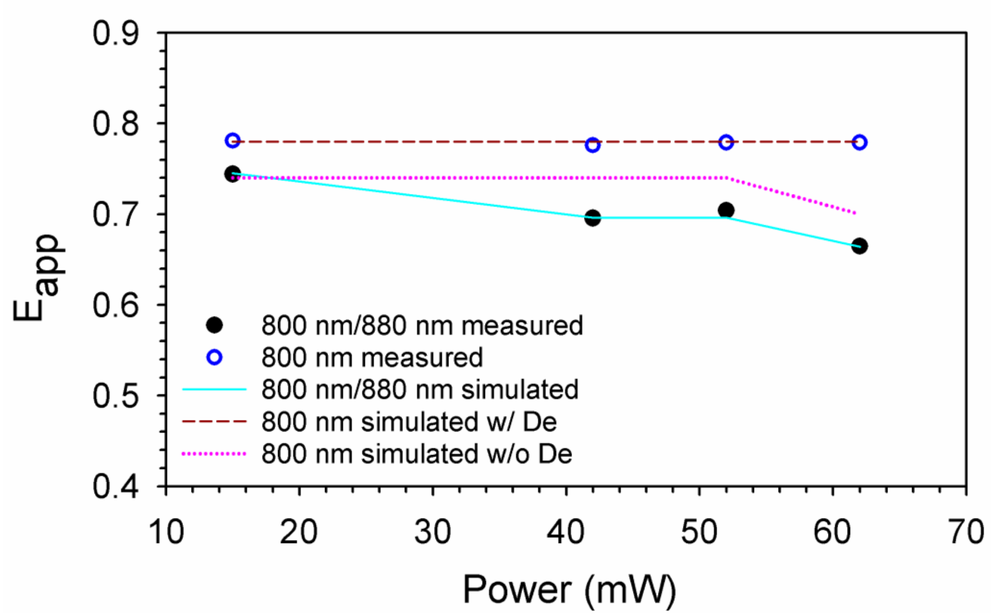


**Supplemental Figure SR3. Comparison of measured and simulated FRET efficiencies (*$E_{app}$*) for the cytoplasmic FRET construct, ADA, obtained using 800 nm and 880 nm excitation at various powers.** The black solid circles represent the weighted average FRET efficiencies across all measurements performed using dual-wavelengths excitation method at 800 nm and 880 nm. Blue empty circles represent the weighted averaged FRET efficiencies across all the measurements for single-wavelength excitation at 800 nm. The cyan solid line represents the best-fit simulated FRET efficiencies for the excitation pair 800 nm/880 nm. Dark red dashed line represents the simulated FRET efficiencies using single excitation at 800 nm with (w/) direct excitation (De) of acceptor. Pink doted line represents the simulated FRET efficiencies using single excitation at 800 nm without (w/o) direct excitation of acceptor.

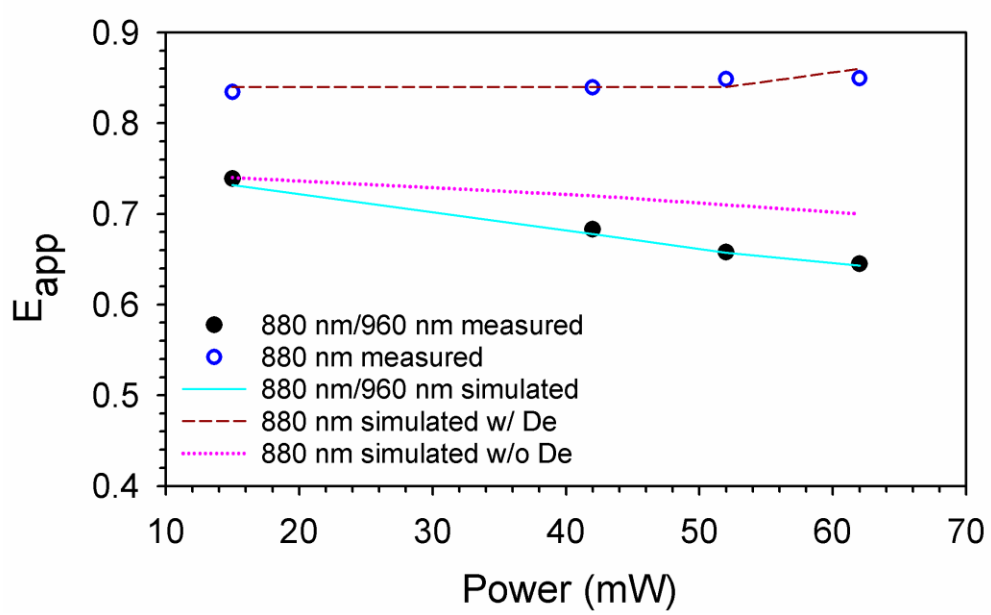


**Supplemental Fig. SR4. Comparison of measured and simulated FRET efficiencies (*$E_{app}$*) for the cytoplasmic FRET construct, ADA, obtained using 880 nm and 960 nm excitation at various powers.** The black solid circles represent the weighted average FRET efficiencies across all measurements performed using dual-wavelengths excitation method at 880 nm and 960 nm. Blue empty circles represent the weighted averaged FRET efficiencies across all the measurements for single-wavelength excitation at 880 nm. Whereas, cyan solid line represents the best-fit simulated FRET efficiencies for the excitation pair 880 nm/960 nm. Dark red dashed line represents the simulated FRET efficiencies using single excitation at 880 nm with (w/) direct excitation (De) of acceptor. Pink doted line represents the simulated FRET efficiencies using single excitation at 880 nm without (w/o) direct excitation of acceptor.
